# Myocardin-related transcription factor drives epithelial fibrogenesis in polycystic kidney disease

**DOI:** 10.1101/2024.03.15.585204

**Authors:** Zsuzsanna Lichner, Mei Ding, Tarang Khare, Qinghong Dan, Raquel Benitez, Mercédesz Praszner, Xuewen Song, Rola Saleeb, Boris Hinz, York Pei, Katalin Szászi, András Kapus

**Affiliations:** Keenan Research Centre for Biomedical Science of St. Michael’s Hospital; Enrich Bioscience, Toronto, ON, Canada; Division of Nephrology, University Health Network, Toronto, ON, Canada; Department of Laboratory Medicine and Pathobiology, Temerty Faculty of Medicine, University of Toronto, Toronto, ON, Canada; Faculty of Dentistry, University of Toronto, Toronto, ON, Canada; Department of Surgery, University of Toronto, Toronto, ON, Canada M5B 1T8; Department of Biochemistry, University of Toronto, Toronto, ON, Canada M5B 1T8

## Abstract

Polycystic kidney disease (PKD) is characterized by extensive cyst formation and progressive fibrosis. However, the molecular mechanisms whereby the loss/loss-of-function of Polycystin 1 or 2 (PC1/2) provokes fibrosis are largely unknown. The small GTPase RhoA has been recently implicated in *cystogenesis*, and we have shown that the RhoA/cytoskeleton/myocardin-related transcription factor (MRTF) pathway is a key mediator of epithelium-induced fibrogenesis. Therefore, we hypothesized that MRTF is activated by PC1/2 loss and plays a critical role in fibrogenic reprogramming of the epithelium. Loss of PC1 or PC2 induced by siRNA in vitro activated RhoA, caused cytoskeletal remodeling and robust nuclear MRTF translocation and overexpression. These phenomena were also manifest in PKD1 (RC/RC) and PKD2 (WS25/-) mice, with MRTF translocation and overexpression occurring predominantly in dilated tubules and in cyst-lining epithelium, respectively. In epithelial cells, a large cohort of PC1/PC2 downregulation-induced genes was MRTF-dependent, including cytoskeletal, integrin-related, and matricellular/fibrogenic proteins. Epithelial MRTF was necessary for paracrine priming of fibroblast-myofibroblast transition. Thus, MRTF is a critical novel mediator of the PC1/2 loss-induced profibrotic epithelial phenotype, and consequently PKD-related fibrosis.

## Introduction

Autosomal dominant polycystic kidney disease (hereafter PKD) is the most common renal genetic disorder with a prevalence of 1:500-1000 (Bergmann et al., 2018; Levy & Feingold, 2000). PKD is caused by loss-of-function mutations in Polycystin 1 (PC1, encoded by *PKD1,* 85%) or Polycystin 2 (PC2, encoded by *PKD2*, 15%). They can form a heteromultimer and act as chemo- and mechanosensitive complex (Douguet, Patel & Honoré, 2019; Luo et al., 2023), impacting a variety of functions, including cell division and metabolism (Kasahara & Inagaki, 2021; Gopalakrishnan et al., 2023; Wang & Dynlacht, 2018). Histologically, the major features of PKD are fluid-filled cysts and progressive renal fibrosis, which ultimately ruin the kidney architecture, leading to End Stage Renal Disease (ESRD) (Fragiadaki, Macleod & Ong, 2020; Zhang, Reif & Wallace, 2020). Intensive work has uncovered fundamental mechanisms in cystogenesis, linking PC1/PC2 loss-of-function to multiple signaling pathways (Caplan, 2022; Chapin & Caplan, 2010; Ferreira, Watanabe & Onuchic, 2015; Márquez-Nogueras, Vuchkovska & Kuo, 2023). However, much less is known about the mechanism underlying the accompanying fibrogenesis.

PKD-related fibrosis is a pathologically important biological puzzle: the disease is epithelium-initiated, but affects the mesenchymal compartment. Recent work, including our own, has shown that epithelial injury is a key causative factor in renal fibrosis (Grande et al., 2015; Bialik et al., 2019; Grgic et al., 2012; Kang et al., 2015; Lovisa et al., 2015; Qiu et al., 2018). Injured epithelial cells undergo partial epithelial-mesenchymal transition, acquiring a profibrotic epithelial phenotype (PEP) (Bialik et al., 2019). This state is characterized by cytoskeletal remodeling and production of profibrotic cytokines/matricellular proteins, which in turn induce the surrounding mesenchyme to become matrix-producing fibroblasts and contractile myofibroblasts (Liu et al., 2020; Zhou et al., 2017; Tan, Zhou & Liu, 2016; Bialik et al., 2019). We and others have shown that MRTF-A, and its isoform MRTF-B (MRTF) are key mediators of the initial epithelial reprogramming and the consequent mesenchymal response (Yamamura et al., 2023; Mao et al., 2020; Speight et al., 2016; Rozycki et al., 2016; Xu et al., 2015; Fintha et al., 2013; Fan et al., 2007; Bialik et al., 2019; Masszi et al., 2010) (reviewed in (Miranda et al., 2021)). Under resting conditions, MRTF resides mostly in the cytosol because it binds G-actin, which masks its nuclear localization sequence (NLS) and promotes its nuclear efflux (Vartiainen et al., 2007; Kuwahara et al., 2005; Miralles et al., 2003). Cellular injury triggers actin polymerization, primarily mediated by the activation of Rho GTPases, such as RhoA. This leads to the dissociation of G-actin from MRTF, and its consequent nuclear translocation. In the nucleus, MRTF binds its cognate transcription factor partner, serum response factor (SRF), and the complex drives a set of genes encoding cytoskeletal, matricellular/matrix proteins and soluble fibrogenic mediators (Esnault et al., 2014; Luchsinger et al., 2011; Small et al., 2010; Morita, Mayanagi & Sobue, 2007; Iwanciw et al., 2003; Mao et al., 2020; Fan et al., 2007; Bialik et al., 2019; Masszi et al., 2010). In addition, SRF can be switched on by an MRTF-independent mechanism, via the ternary complex factors (TCFs) (Gualdrini et al., 2016), leading to the activation of immediate early genes (c-Fos, c-Myc) regulating cell cycle entry and proliferation, relevant for PKD pathogenesis (Lakhia et al., 2023).

Remarkably, enhanced RhoA signaling has been recently implicated in PKD as an important input for *cystogenesis*. PC1 loss-of-function was shown to induce RhoA activation (Streets, Prosseda & Ong, 2020; Cai et al., 2018) which in turn triggered Rho kinase- and myosin phosphorylation-dependent activation of the Hippo pathway effector YAP and its paralog TAZ, facilitating cyst formation. c-Myc was identified as a critical YAP target gene driving *cystogenesis* (Cai et al., 2018). This scenario raised the intriguing possibility that MRTF, downstream of activated RhoA signaling, might mediate the other major arm: epithelial fibrogenesis. In further support of this hypothesis, a subset of MRTF and YAP target genes are common (Foster, Gualdrini & Treisman, 2017; Kim et al., 2017; Yu, Miyamoto & Brown, 2016; Speight et al., 2016; Bialik et al., 2019), both MRTF and YAP/TAZ contribute to PEP (Speight et al., 2013; Rozycki et al., 2016; Bialik et al., 2019) and MRTF itself is an upstream transcription factor for TAZ (Liu et al., 2016; Miranda et al., 2017). Further, a systems biology approach has identified SRF as a major dysregulated transcription factor pathway in PKD (Song et al., 2009a). However, it remained to be elucidated whether MRTF is activated/translocated upon PC1/2 loss or in PKD, and whether it is a significant contributor to a fibrogenic signature. Accordingly, our goal was to place MRTF on the “PKD map” and to link it to fibrogenesis.

Our results show that absent/dysregulated PC1 and PC2 signaling induces nuclear translocation of MRTF in tubular epithelial cells in a RhoA-dependent manner. MRTF in turn facilitates the epithelial expression of cytoskeletal components, integrins, matricellular proteins and other mediators, thereby priming mesenchymal cells for fibrogenesis.

## Results

### PC1 or PC2 downregulation activates RhoA and induces nuclear translocation of MRTF

To test whether PC1 or PC2 downregulation affected MRTF signaling, we silenced the corresponding genes in LLC-PK1 tubular cells, using specific siRNAs. This approach efficiently and selectively reduced the expression of the corresponding proteins allowing their independent manipulation (Figure 1A, Expanded View Figure 1). To assess whether these proteins impact RhoA activity in our cellular system, we measured the level of active (GTP-bound) RhoA using GST-Rhotekin pull-down assay, as previously (Kakiashvili et al., 2009). Active RhoA (normalized to total RhoA) was significantly elevated upon PC1 loss (Cai et al., 2018; Streets, Prosseda and Ong, 2020), and similar results were obtained for PC2 (Figure 1B). Quantitative immunofluorescence staining with an active RhoA-specific antibody corroborated these results (Figure 1C). Loss of PC1 or PC2 resulted in dramatic cytoskeletal remodeling, characterized by the formation of strong actin stress fibers (Figures 1D). This was accompanied by a substantial nuclear translocation of MRTF. Both of these changes were prevented/mitigated by concomitant downregulation of RhoA (Figure 1D). These qualitative observations were quantified by two means. First by visual inspection, using a tripartite compartmental distribution (cytosolic, nuclear, both/even) of MRTF. While MRTF was predominantly cytosolic under control conditions, it exhibited a significant shift to the nucleus upon loss of PC1 or PC2. This was reversed by RhoA downregulation (Figure 1E). Second, to overcome regional heterogeneity, the distribution of single-cell nuclear/cytosolic MRTF ratios were automatically determined in large cell populations (detailed in Methods). PC1 or PC2 silencing resulted in a shift toward higher N/C ratios (black vs. red curves), which was strongly mitigated by RhoA silencing (Figure 1F). The cumulative frequency of N/C ratios ≥1.4 (univocally assessed as “nuclear MRTF” by visual inspection) showed a ≈6-fold rise in PC1- or PC2-silenced cells relative to controls; this change was abolished by RhoA downregulation (Figure 1G). Together these results show that the loss of PC1 or PC2 induces robust RhoA-dependent MRTF translocation into the nucleus.

**Figure 1.**
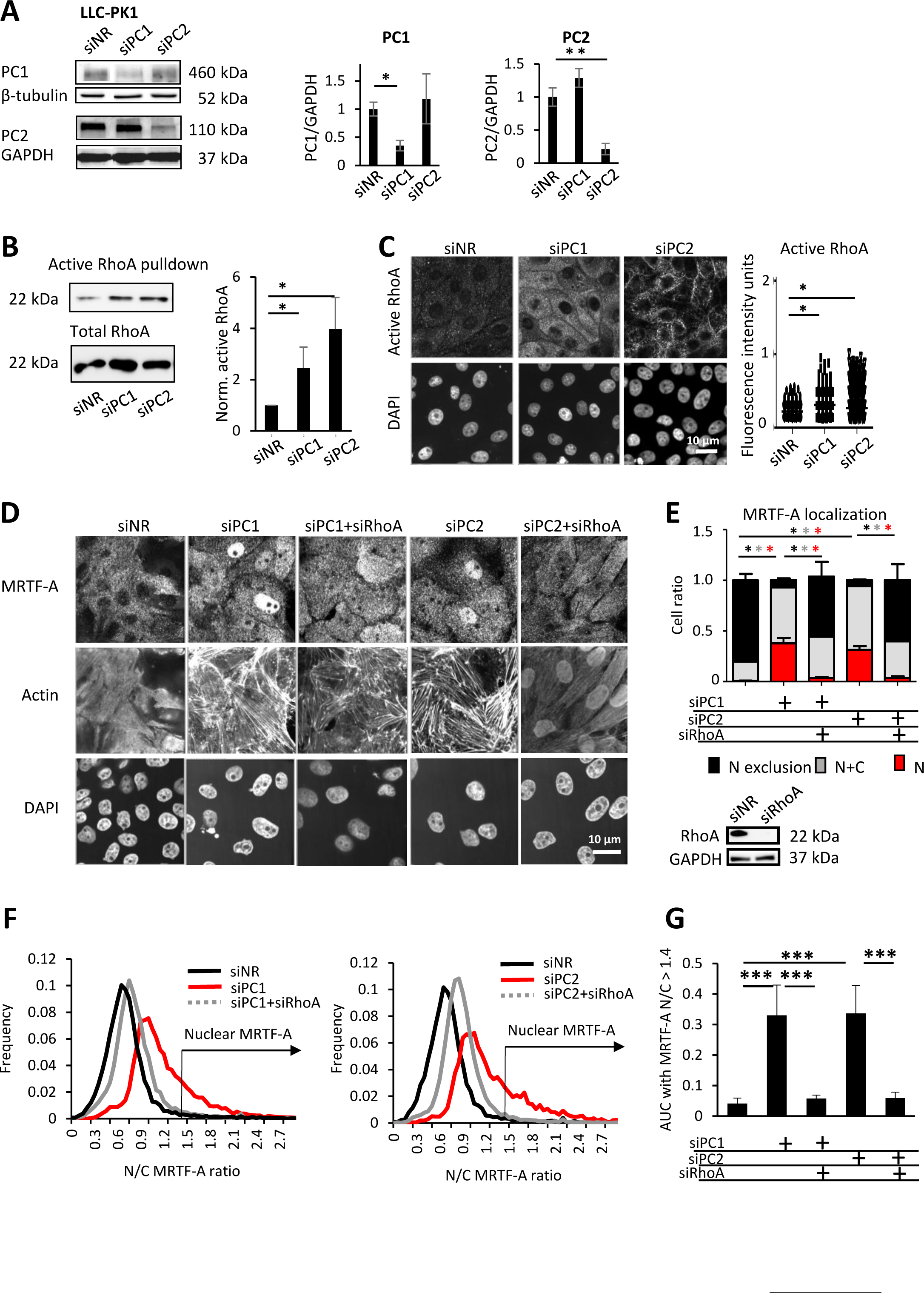
Loss of Polycystin 1 or 2 activate RhoA and leads to RhoA-dependent cytoskeletal remodeling and nuclear accumulation of MRTF-A. (A) PC1 or PC2 were silenced in LLC-PK1 cells with the corresponding siRNAs (150 nM and 100 nM, respectively, 48 h), without altering each other’s expression, as assessed by Western blot analysis. PC/GAPDH ratios are presented in the right panel. (B) Active RhoA was detected by GST-Rhotekin binding domain precipitation assay. Active RhoA precipitates and total cell lysate blots were probed with a RhoA-specific antibody. Fold change was normalized against the active RhoA/total RhoA ratio of control siRNA-transfected cells (non-related siRNA or siNR). (C) Cells were transfected as above. After 48 h, active RhoA was detected by immunofluorescence, using an antibody specific for the active, GTP-bound form. Representative images are shown for each indicated condition. Fluorescence intensity was quantified in individual cells using ImageJ. (D, E) PC1 (150 nM siRNA) or PC2 (100 nM siRNA) were silenced alone or concomitantly with RhoA (50 nM siRNA), as indicated. MRTF-A subcellular localization was assessed by immunofluorescent staining in each condition. In parallel, F-actin was visualized by staining with iFLUOR555-labelled phalloidin (representative images of the brightest plane are shown in D). Note that RhoA knockdown was highly efficient (E). Using visual assessment of MRTF-A subcellular localization, cells were grouped into the following three categories: 1. predominantly nuclear MRTF-A (red bars, N=nuclear), 2. even nuclear and cytoplasmic presence of MRTF-A (grey bars, N+C), 3. nuclear exclusion of MRTF-A (black bars, N exclusion). * indicates p<0.05, each colored * indicates the significance of the corresponding bar. (F) MRTF-A subcellular localization was analyzed by ImageXpress, a high-throughput automated digital imaging system. The distribution curves indicate the frequency of cells with increasing nuclear-to-cytoplasmic MRTF-A ratio (N/C). (G) Cumulative quantitation of the distribution curves shown in F, using N/C > 1.4 as a cutoff, which corresponds to definite nuclear MRTF-A localization by visual classification.

### PC1/2 loss elevates MRTF expression

During these experiments, we noted that PC loss affected not only the distribution, but also the total expression of MRTF. Indeed, MRTF-A immunostaining significantly increased both in PC1- and PC2-silenced cells (Figure 2A and 2B), a finding confirmed by Western blots (Figure 2C). This rise was, at least in part, due to increased MRTF gene transcription, since PC1 and PC2 downregulation significantly elevated the message for both MRTF-A and MRTF-B (Figure 2D). These findings imply that PC1/2 loss facilitates MRTF signaling both at the level of activation/nuclear translocation and of total expression.

**Figure 2.**
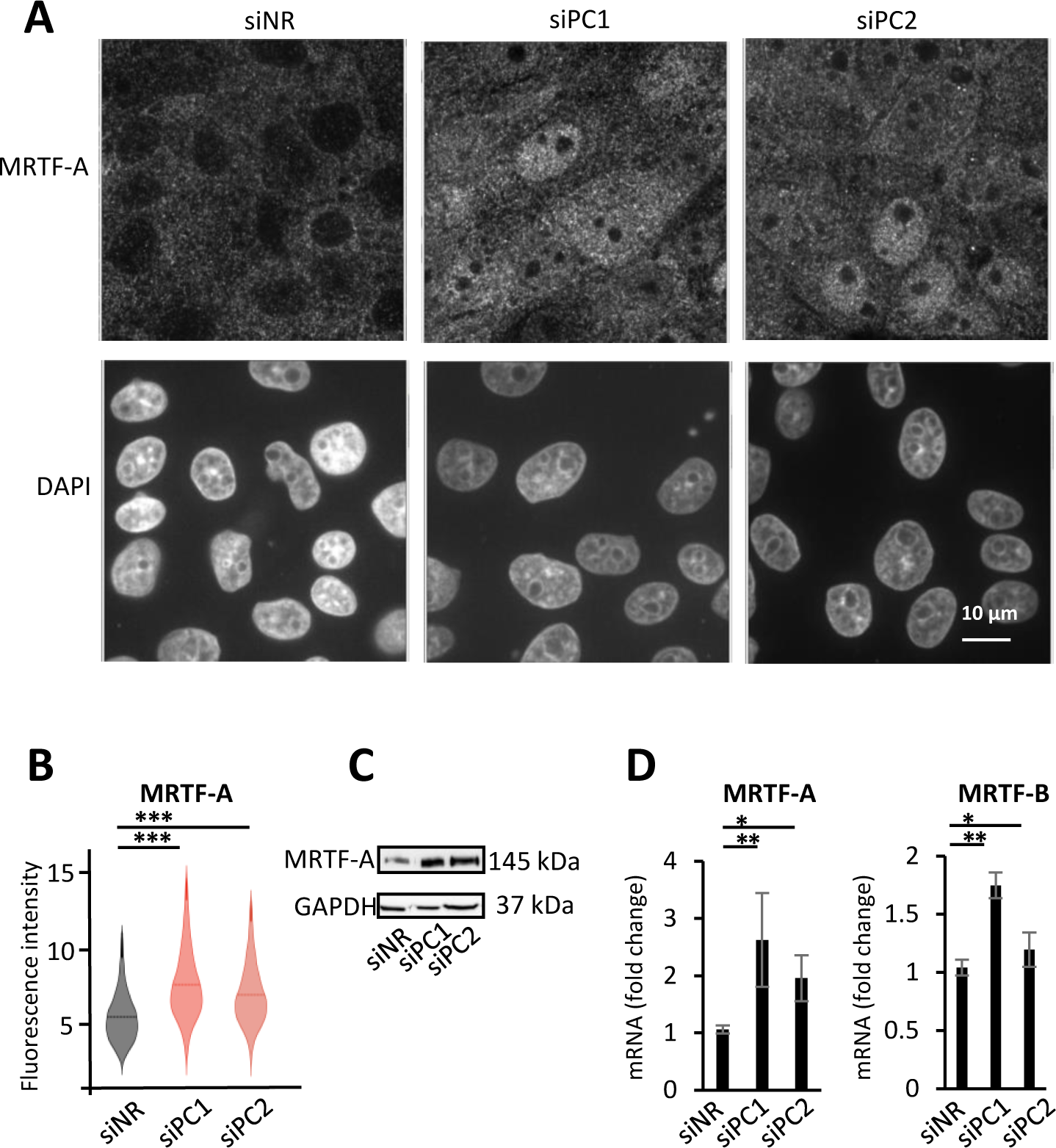
MRTF-A expression is upregulated by Polycystin loss. PC1 and PC2 were downregulated by siRNA transfection (150 nM and 100 nM, respectively, 48 h) and MRTF-A expression was assessed by three methods. (A, B) MRTF-A-specific fluorescence intensity was quantified by ImageXpress (see Materials and Methods). Violin plots indicate the median fluorescence value. (C) MRTF-A protein expression was also visualized by Western blot analysis. (D) Both MRTF-A and MRTF-B mRNA expression was quantified by RT-qPCR. Results were normalized to PPIA. * p<0.05, **, p<0.01, *** p<0.001.

### MRTF in action upon PC loss in the epithelium: a targeted approach

Next, we addressed the functional significance of the observed changes. As an initial strategy, we concentrated on genes that satisfied each of the following criteria: they were shown to be a) PC-sensitive (Lanktree et al., 2021; Yook et al., 2012; Song et al., 2009a) b) MRTF targets (Esnault et al., 2014) and c) involved in fibrogenesis/PEP. Downregulation of PC1 (Figure 3A, left panels) or PC2 (Fig 3A right panels) significantly enhanced the mRNA expression for matricellular proteins/fibrogenic mediators, such as transgelin (TAGLN) (see also Expanded View Figure 1 for alternative PC1 and PC2 siRNAs), CTGF and CYR1. Downregulation of MRTF-A strongly reduced TAGLN and CTGF expression upon PC1 or PC2 silencing, and significantly decreased CYR61 expression in PC1-but not in PC2-downregulated cells (Figure 3A). Efficient MRTF-A downregulation was not altered by concomitant silencing of PC1 or PC2 (Expanded View Figure 1B). The impact of MRTF-A was preserved when PC1/2- downregulated cells were exposed to TGFβ1, the most potent fibrogenic cytokine, which is elevated in and promotes PKD-associated fibrosis (Hassane et al., 2010; Zhang et al., 2020). TGFβ1 potentiated/augmented the effect of PC1/PC2 loss for each of these three genes, and MRTF significantly reduced the combined effect of these stimuli, except for PC2 downregulation-induced CYR61 expression (Figure 3A). PC1 or PC2 loss also stimulated the expression of ANXA1 and RASSF2 in an MRTF-A-dependent manner (Figure 3B), although these were not stimulated by TGFβ1 (not shown). We used the most responsive gene, TAGLN, to test RhoA- and SRF-dependence as well. Downregulation of either efficiently reduced PC1/2 loss-provoked TAGLN expression (Figure 3C, 3D). MRTF-B also contributed to these responses, because its silencing also mitigated the PC1- or PC2 loss-triggered increase of TAGLN, CTGF and CYR61. Interestingly, in PC2-downegulated cells CYR61 mRNA expression was selectively sensitive to MTRF-B depletion (Figure 3E-F).

**Figure 3.**
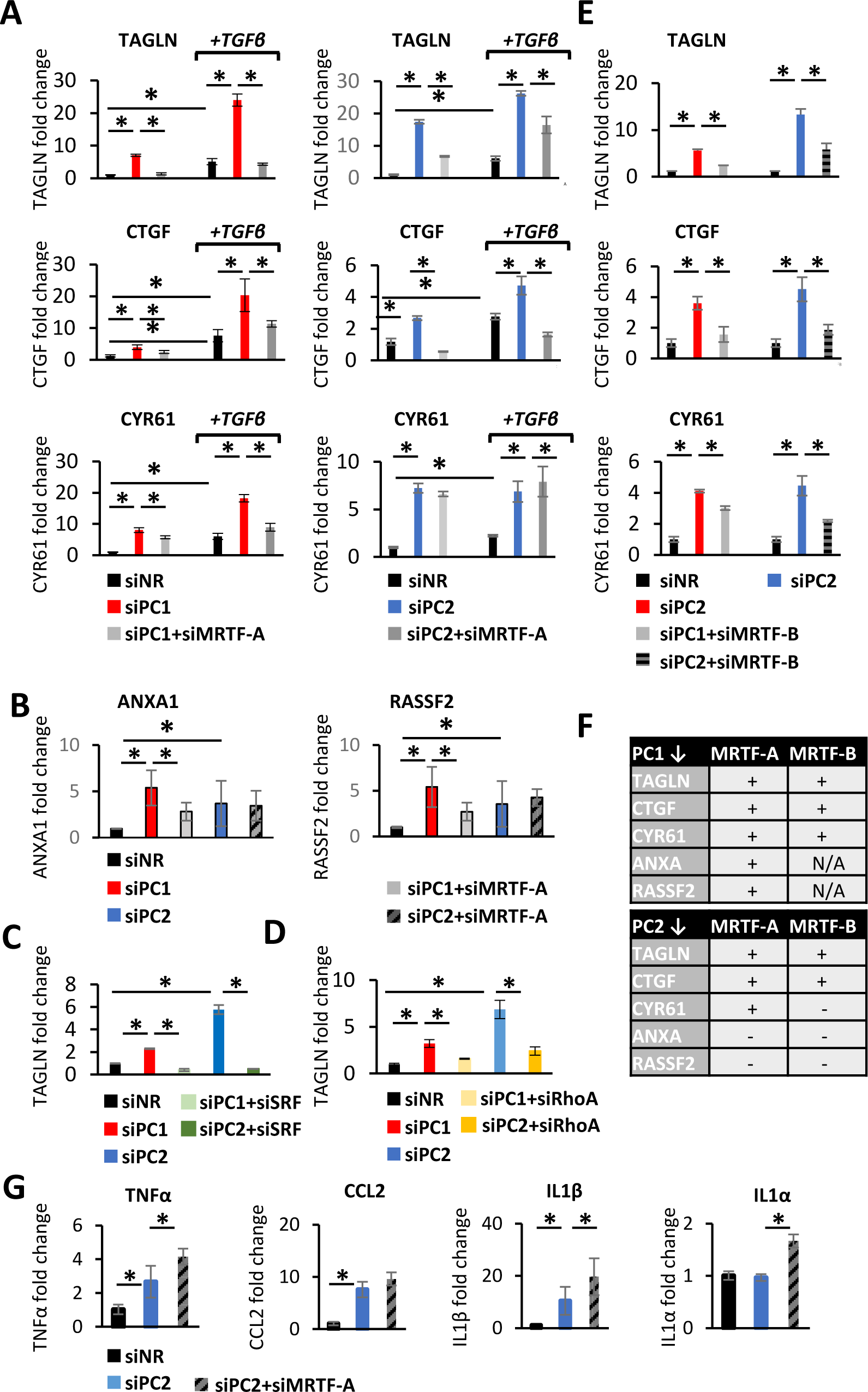
Polycystin loss induces the expression of profibrotic phenotype-related transcription program in a partially MRTF-dependent manner. (A-E, G) LLC-PK1 cells were transfected with control siRNA (NR) or siRNAs targeting PC1 (150 nM), PC2 (100 nM), MRTF-A (50 nM), MRTF-B (50 nM), SRF (100nM) or RhoA (100nM), as indicated. In (A, B), 24 h post-transfection, cells were treated with DMSO or 5 ng/ml TGFβ in serum-free media for an additional 24 h. Synergy/additional effects between the loss of PCs and the presence of TGFβ was assessed by measuring the expression of known PEP-related genes that are direct transcriptional targets of MRTF-A. The synergistic effects varied among genes, as shown. ANXA and RASSF expression was not stimulated by TGFβ treatment, but was stimulated by PC loss and was partially MRTF-driven. Fold changes (normalized to PPIA expression) were compared to the DMSO-treated, siNR-transfected samples (n>4 in triplicates). (C-D) Knockdown of RhoA and SRF diminished the profibrotic effect of PC loss. (F) The table summarizes the MRTF-dependence of the investigated PEP-related and validated MRTF/SRF targets. (G) Silencing MRTF-A facilitates the PC loss-induced increase of some Nuclear factor κB (NFκB)-dependent inflammatory genes. mRNA abundance for tumor necrosis factor-α (TNFα), chemokine (C-C motif) ligand 2 (CCL2), interleukin-1β and α (IL**-**1β, IL-1α) are shown. * p<0.05

Next, we quantified mRNA expression for some pro-inflammatory genes, cognizant that MRTF can physically associate with and inhibit NFκB (Hayashi et al., 2015; Wang et al., 2012a). PC2 downregulation stimulated TNFα, CCL2, and IL1β expression, while did not alter IL1α mRNA levels. MRTF-A silencing significantly *potentiated* the effect of PC2 loss on TNFα and IL1β and increased IL1α mRNA expression (Figure 3G). Thus, MRTF is an important contributor of PC loss-induced expression of key fibrogenic/PEP genes (Figure 3F), and a suppressor of some proinflammatory genes, shifting the balance from acute epithelial inflammatory responses to fibrosis.

### MRTF in action upon PC loss: a general approach

To assess the role of MRTF in the molecular pathogenesis of PKD from a wider angle, we performed RNA-Seq. We focused on genes that a) were upregulated upon PC1 *or* PC2 silencing *and* b) were MRTF-dependent. Three complementary analysis methods were used to identify such PKD-related and MRTF-dependent events (Figure 4A).

**Figure 4.**
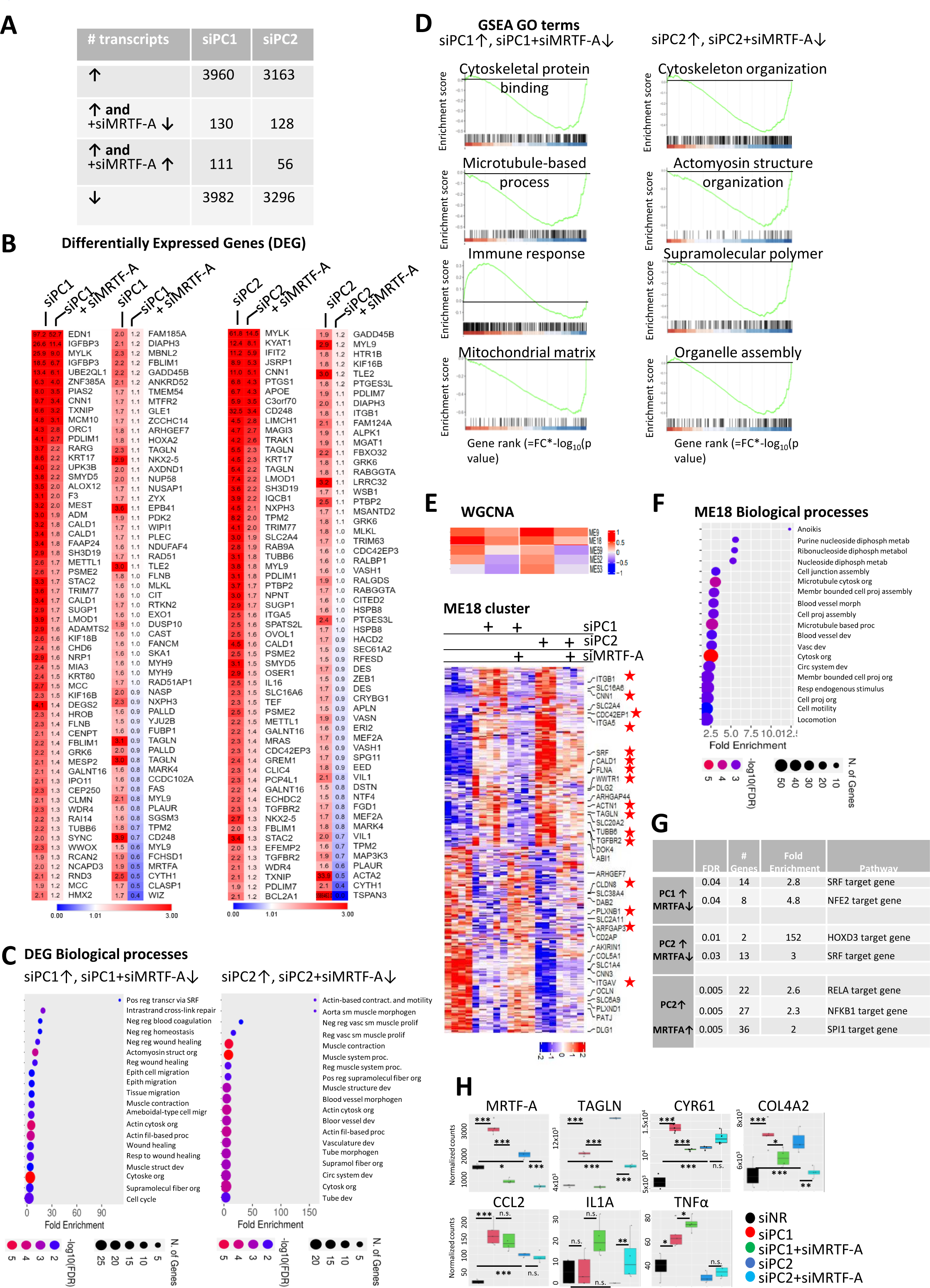
Next generation transcriptome analysis of Polycystin- and MRTF-A-dependent gene expression. (A) Overview of experimental framework. LLCPK-1 cells were transfected with control siNRA (siNR, 200nM, n=3), siPC1 (150 nM, n=4), siPC2 (100 nM, n=4), alone or with siMRTF-A (100nM, siPC1+siMRTF-A n=5, siPC2 n=4). Gene expression was compared across the indicated conditions using RNA-Seq. 1.5 fold cutoff defined upregulation and downregulation (log_2_ >0.6 upregulated, log_2_<-0.6 downregulated, p≤0.05, base mean ≥50). The table indicates the number of transcripts upregulated upon PC knockown (↑), and significantly changed when MRTF was also silenced. (B) DEG Analysis. Expression heatmap of transcripts that are upregulated by PC1 or PC2 loss (siPC1 and siPC2) in an MRTF-dependent manner (siPC1+siMRTF-A and siPC2+siMRTF-A). Numbers indicate fold changes compared to non-related siNR-transfected LLC-PK1 cells (siNR). (C) Shinygo 0.76 (a web application for R) was used to search for GO Biological Processes that are enriched when PC expression is silenced and are MRTF-dependent. (D) Gene Set Enrichment Analysis (GSEA) defined enriched GO terms in the above-described comparisons, as indicated. Selected significant GO terms are shown. (E) Expression heatmap of significant Weighted Gene Co-expression Analysis (WGCNA) modules that were MRTF-dependent and upregulated in PC1 or PC2 knockdown conditions. Heatmap of ME18 module is presented separately in the lower panel. Stars indicate genes that are known direct targets of MRTF. (F) The graph is the visual representation of significant GO biological processes related to ME18 module. (G) Predicted key transcription factors for genes that were upregulated upon PC knockdown and showed expression change upon MRTF-A silencing (Transfac TF Binding Site Enrichment Analysis). (H) Expression profiles from the RNA-Seq data shown for some individual genes under various conditions, as indicated. Key PKD-related genes (MRTF-A, TAGLN, CYR61, COL4A2) and innate immunity-related genes (lower panels) are shown. Adjusted p-values are indicated: * p<0.05, ** p<0.01, *** p<0.001.

First, we identified *differentially expressed genes/transcripts* (DEG) that were significantly elevated upon PC1 or PC2 downregulation *and* significantly suppressed by concomitant MRTF silencing. The corresponding 130 (PC1/MRTF) and 128 (PC2/MRTF) transcripts are shown in (Figure 4B). Common enriched biological processes included epithelial cell migration, wound healing, and cell contractility, in line with the key role of MRTF in early epithelial injury responses (Figure 4C, Expanded View Figure 2-4).

Second, *Gene Set Enrichment Analysis* (GSEA) (Figure 4D) indicated that the actomyosin cytoskeleton was among the most significant MRTF-dependent PKD-associated GO terms. The presence of closely related categories (microtubules, supramolecular polymers) underlines the relationship between PKD, cytoskeletal reorganization and MRTF. GSEA also identified ‘mitochondrial matrix’ and ‘organelle assembly’ as MRTF-supported categories. The MRTF-dependence of these is of special interest as altered cellular metabolism is a hallmark of PKD (Podrini, Cassina & Boletta, 2020; Padovano et al., 2018). Moreover, MRTF emerged as a significant negative regulator of the early immune response.

Third, we utilized *Weighted Gene Co-expression Network Analysis (WGCNA*), to identify unbiased positive and negative correlation patterns. Among the resulting 90 modules, six matched our expression criteria, supplemented with the extra-requirement that PC1 *and* PC2 loss should act concordantly. Of these, ME18 was ranked first and it was the second most significant among all the 90 modules (Figure 4E). ME18-related biological processes reflected the previously identified themes (cell junction, microtubules, cytoskeletal organization, cell projection assembly, cell motility) (Figure 4F).

Considering common cis-elements in the target genes, SRF-binding sites were highly enriched in the MRTF-dependent, PKD-related gene promoters, as expected. Interestingly, the recognition elements of innate immune response-related transcription factors (particularly NFκB) were also significantly enriched, concordant with the DEG analysis, suggesting that MRTF inhibits NFκB (Figure 4G-H).

Finally, we confirmed the PC1/2 status-independent downregulation of MRTF-A by siRNA and the concomitant suppression of the PC1/2 loss-promoted TAGLN and the PC1-specific response of CYR61 (Leask, 2013; Lee et al., 2015; Doke et al., 2022). In addition, we also show the MRTF-dependent behavior of Col4A2, a basement membrane component, the expression of which was shown to increase in renal fibrosis (Bialik et al., 2019) (Figure 4H). Thus, RNA-Seq indicates that MRTF significantly contributes to PKD-associated transcriptome alterations in our epithelial cell model system, by enhancing cytoskeletal reorganization and matricellular proteins, which are key features of PEP. In addition, it regulates mitochondrial organization and metabolism and mitigates the acute innate immune response.

*Integrin β1 (ITGB1) and MRTF-A form a feed-forward loop and regulate profibrotic gene expression* Our transcriptome sequencing data also indicated that PC loss stimulated the expression of ITGB1 (Figure 5A). Relevantly, deletion of ITGB1 dramatically reduced cystogenesis and fibrosis in PKD1 fl/fl, Aqp2-Cre animals (Lee et al., 2015). Further, ITGB1 is a known direct MRTF target, whose promoter harbors a CARG box (Brandt et al., 2009b, 2009a). Using qPCR, we verified that PC1 and PC2 knockdown increased ITGB1 mRNA expression. This transcriptional change was partially MRTF-A-dependent (Figure 5B). Moreover, ITGB1 *activity* was also affected by PC loss; using an activation-specific ITGB1 antibody (12G10), we observed a major increase in the number of active ITGB1 clusters, visualized as parallel thin lines in PC1- or PC2-silenced cells. Similar effect was observed upon the addition of manganese (Mn^2+^), a pan-integrin activator, which was used as a positive control (Figure 5C) (Ni et al., 1998). Importantly, Mn^2+^ treatment alone was sufficient to prompt MRTF-A nuclear accumulation and robust TAGLN expression (Figure 5C-F). To assess the potential contribution of ITGB1 to MRTF translocation, we treated the cells with ITGB1 siRNA, which resulted in ≈70% drop in ITGB1 mRNA (Figure 5E left panel).

**Figure 5.**
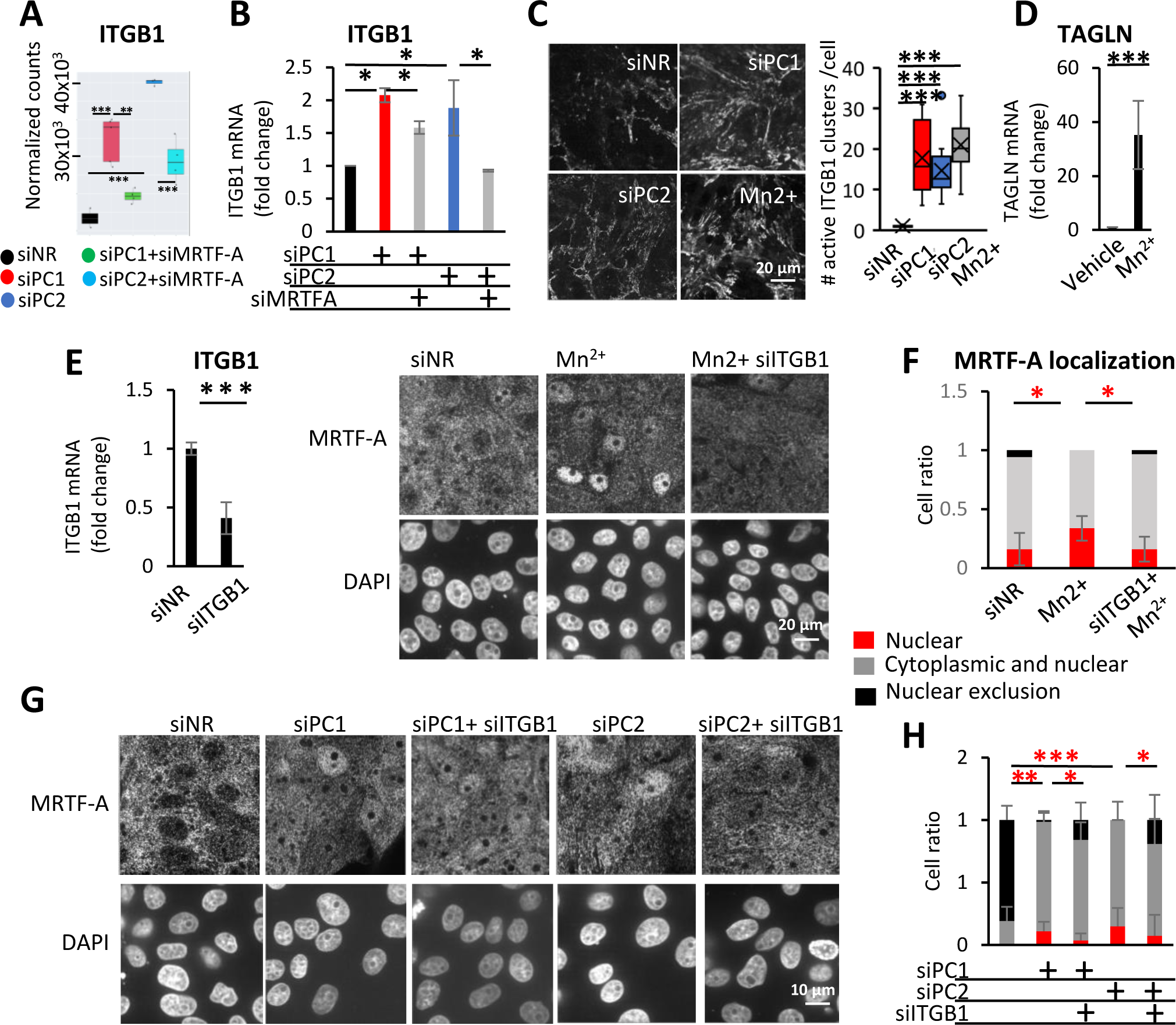
ITGB1 is overexpressed and activated upon Polycystin loss and regulates the subcellular localization of MRTF. (A, B) ITGB1 mRNA expression was quantified using RNA-Seq and RT-qPCR in LLC-PK1 cells transfected with siPC (siPC1 150 nM or siPC2 100 nM) or siNR, and siMRTF-A (50 nM) or siNR, as indicated. RT-qPCR results were normalized to PPIA expression, against the siNR-transfected controls. (C) Left panel: Antibody, specific for the active conformation of ITGB1 (12G10) was used to visualize the clustering of active ITGB1. Right panel presents the number of ITGB1 clusters, normalized to cell number. (D) Mn^2+^ treatment (500 μM, 1 h) activated integrins and was sufficient to drive high TAGLN expression. Relative expression was measured by qRT-PCR and was normalized against PPIA expression. (E) ITGB1 was partially silenced by specific siRNA (100 nM, 48 h). (F) Integrins were activated by Mn^2+^ treatment and MRTF-A subcellular localization was estimated by immunofluorescence staining. Using visual assessment (25 fields), MRTF-A localization was categorized as nuclear (red bars), even (grey bars) or cytoplasmic (black bars) (right panel). (G, H) Cells were transfected with PC1 (150 nM) or PC2 (100 nM) alone or together with siNR or siITGB1 (100nM). Representative images of anti-MRTF-A immunofluorescent staining are shown (left panel), and reveal a modest but significant cytoplasmic shift upon ITGB1 silencing. MRTF-A localization was quantified as in D (right panel). The images and quantification represent the results of minimum three independent experiments.

This treatment efficiently suppressed the Mn^2+^-induced nuclear accumulation of MRTF (Figure 5E, F), and significantly, but modestly mitigated PC1 or PC2 loss-provoked MRTF translocation (Figure 5G, H). Together, these finding imply that PC1/2 loss elevates the level and activity of ITGB1; and integrin activation is sufficient to induce MRTF translocation, which could contribute to (but is likely not indispensable for) this effect. Thus, while ITGB1 and MRTF can mutually activate each other, MRTF can also be triggered by other pathway(s) downstream of PC loss (see Discussion).

### The role of MRTF in paracrine epithelial-mesenchymal communication upon PC loss

To access whether PC loss can elicit a functional PEP state inducing fibroblast-MyoF transition (Grande et al., 2015; Bialik et al., 2019; Lovisa et al., 2015; Yang et al., 2018; Yao et al., 2021), and to test whether this might occur in an MRTF-dependent manner, we employed a bioassay. Conditioned media derived from control or siPC1-transfected epithelial cells were transferred onto fibroblasts, and the ensuing fibroblast-MyoF transition was determined based on the capacity of MyoF to contract and wrinkle the underlying soft substrate (Schuster et al., 2023; Wipff et al., 2009) (Figure 6A). Conditioned media of the siPC1-transfected cells induced strong wrinkling capacity in fibroblasts. Importantly, conditioned media derived from cells exposed to simultaneous knockdown of MRTF-A and PC1 lacked this effect (Figure 6B). Thus, PC loss induces the profibrotic paracrine PEP state in an MRTF-dependent manner.

**Figure 6.**
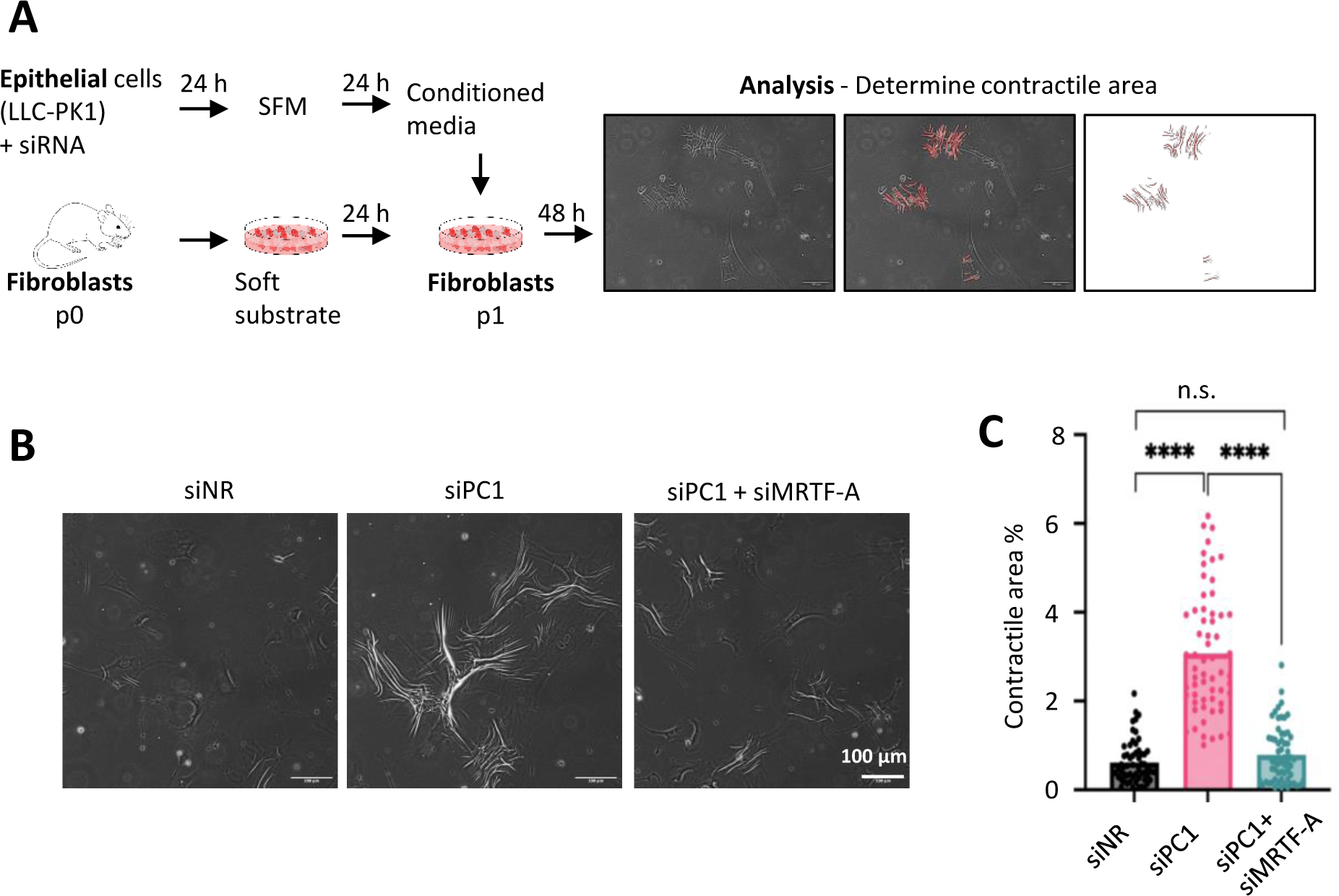
Epithelial Polycystin 1 loss induces MRTF-dependent paracrine signaling that potentiates fibroblast-to-myofibroblast transition. (A) Experimental design. LLC-PK1 cells were transfected with the indicated siRNAs (150 nM siPC1 and 50 nM siNR or siMRTF-A). Twenty-four h post-transfection, cells were thoroughly washed to avoid the carry-over of siRNAs to fibroblast culture, and fresh serum-free media was added. Conditioned media was collected after an additional 24 h and was used to stimulate fibroblasts. Forty-eight h later, >10 randomly selected areas of the fibroblast cultures were photographed under each condition. Cell contractile function, related to % area covered by wrinkles, was analyzed by FIJI ImageJ software and the values were normalized to cell number. (B-C) Representative images of fibroblasts cultures at the experimental end points and their quantification.

### The MRTF pathway is activated in vivo in various forms of PKD

To assess whether PKD affects MRTF distribution *in vivo*, we analyzed two established adult-onset mouse models, PKD2 WS25/- and PKD1 RC/RC (Wu et al., 1998; Hopp et al., 2012; Cai et al., 2018) (Figure 7A). *Pkd2* WS25/- is a compound heterozygous model, wherein the somatically unstable WS25 allele undergoes high rates of recombination resulting in the loss of *Pkd2,* leading to early cystogenesis (within 3 months) (Wu et al., 1998). *PKD1* p.R3277C (RC) is a functional hypomorphic mutation, causing cystogenesis and fibrosis at 12 months of age. Because the younger (3 months) PKD2 WS25/- cohort showed more concordant results, this model was analyzed in more detail. Active RhoA levels were significantly higher PKD2 WS25/- kidneys compared to their corresponding controls (Figure 7B). As expected, PKD2 WS25/- kidney were histologically characterized a large number of dilated tubular structures and cysts (Figure 7C, upper panes). Remarkably, robust nuclear MRTF accumulation was observed in a subset of epithelial cells, predominantly in dilated, precystic tubules (Figure 7C). Nuclear MRTF accumulation appeared to be much less and more homogenous in control animals (Figure 7C). It is worth noting, however, that pathological tubular structures in PKD animals were interspersed among normal ones (with low nuclear MRTF staining), reflecting inherent variance in PKD1 dosage and PKD2 genomic rearrangements, as suggested before (Wu et al., 1998; Hopp et al., 2012). In addition to tubular MRTF accumulation, there was a striking, general (cytosolic and nuclear) upregulation of total cellular MRTF-A expression in the cyst-lining epithelium (Figure 7C, Expanded View Figure 5A). Concordant with these qualitative observations, both total cortical (Figure 7D) and cystic (Figure 7E) MRTF expression was significantly higher in PKD2 kidneys than in the corresponding controls, and it was much higher in cysts than in the normal regions of PKD2 kidneys (Figure 7E). Similar observations (i.e. marked nuclear MRTF accumulation in the tubules and overexpression in the cyst) were made in PKD1 animals as well (Figure 7C, lower panes, 7D). In accordance with these results, PKD patients’ samples also exhibited strong MRTF-A staining in the cystic epithelium (Expanded View Figure 6).

**Figure 7.**
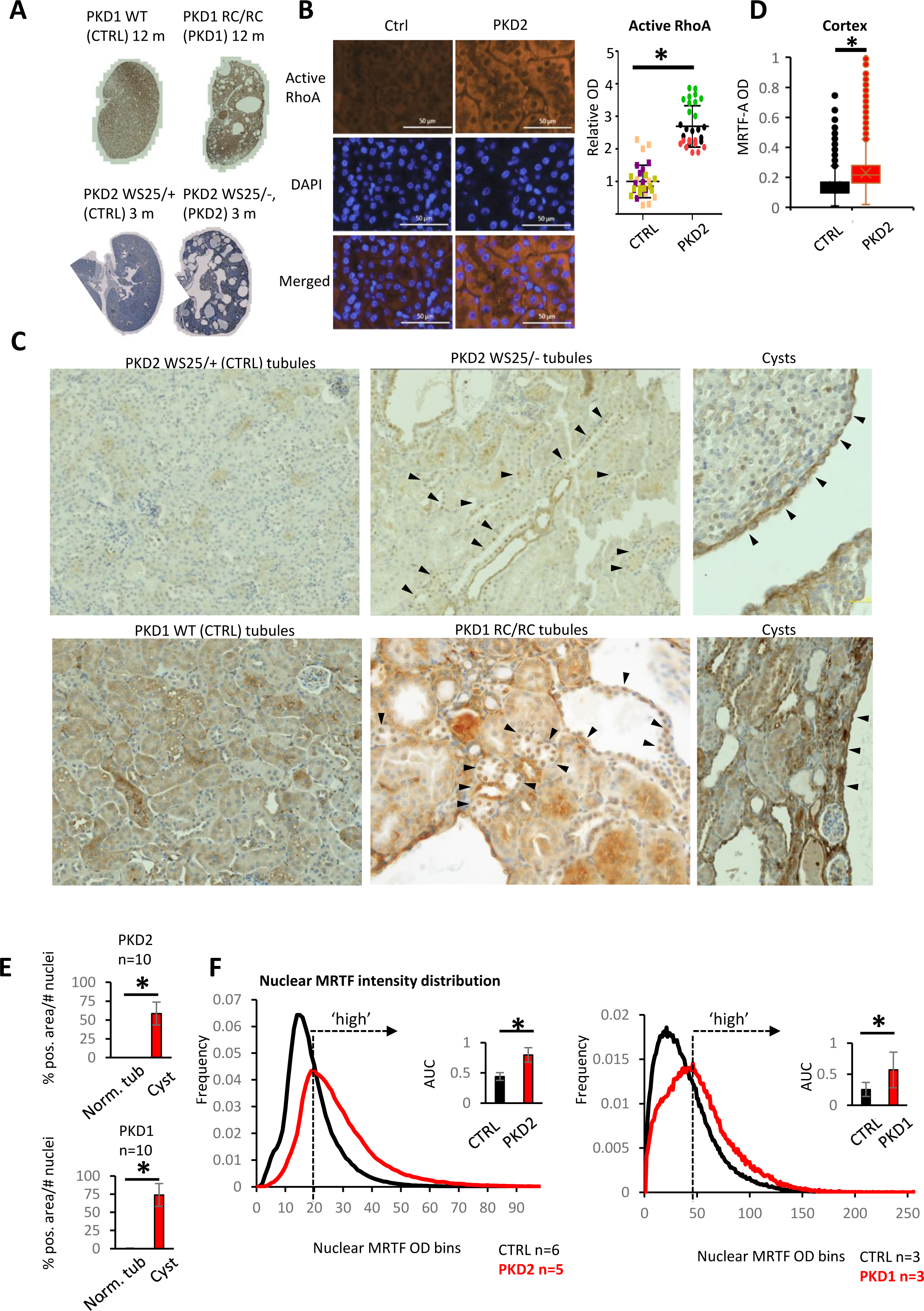
Transgenic PKD mouse model kidneys show MRTF nuclear accumulation and overexpression. (A) Twelve months old PKD1 RC/RC mice (PKD1, n=3), WT control littermates (Ctrl, n=3), 3 months old PKD2 WS25/- animals (PKD2, n=6) and PKD2 WS25/+ control littermates (Ctrl, n=6) were evaluated. At the selected time points micro- and macrocysts were present in both models, except for one of the PKD2 WS25/- mice. This animal was omitted from the analysis. (B) RhoA activation was assessed by detecting the GTP-bound form by immunofluorescence. Ten fields were quantified from each animal (n=3, right panel). (C) Nuclear MRTF expression was quantified in the using HALO’s area quantification package. (D) IHC assessed MRTF subcellular localization in both PKD mouse models and control animals. Arrowheads indicate nuclear MRTF-A in the tubules and increased MRTF-A expression in the cystic epithelial wall. (E) Whole kidney sections were analyzed, MRTF-A/DAB average OD was individually reported for the >300,000 nuclei per animal, using the Object Quantification package from HALO. OD values were assigned to 256 incremental bins, and single nuclei-associated OD data are presented as distribution curves. The highest frequency OD bin in the PKD animals was assigned as the threshold for ‘high’ nuclear MRTF-A accumulation (dotted arrows). The inserts depict the AUC ratio corresponding to nuclei with ‘high’ nuclear MRTF-A. (F) MRTF-A expression on histologically ‘normal’ tubules and cysts were compared within the PKD2 WS25/- or PKD1 RC/RC kidneys. Automatic Threshold Method (ATM score 0 or 1, Area Quantification package, HALO) was used to dichotomize DAB positive and negative pixels. MRTF-A positive area was normalized to the number of nuclei in each examined tubule and cyst.

Considering that kidneys show focal heterogeneity in nuclear MRTF accumulation, we sought to compare this parameter quantitatively on whole kidney sections of normal, PKD2 and PKD1 animals. Therefore, the distribution of nuclear MRTF-A (averaged for all animals tested in whole kidney slices, >300,000 cells/kidney) was measured by the HALO image analysis platform. A substantial shift towards higher nuclear intensities (OD) was observed both in PKD2 and in PKD1 kidneys, compared to the corresponding healthy controls (Figure 7F, Expanded View Figure 5B and 7). The OD corresponding to the maximum of the distribution curve in PKD mutant animals was selected as a threshold defining ‘high’ MRTF-A nuclear presence. As shown in the insets, significantly larger percentage of cells exhibited ‘high’ nuclear MRTF-A expression in the PKD mutant kidneys than in the controls (Figure 7F).

### CTGF expression shows spatial correlation with increased MRTF expression

Under physiological conditions, only stromal cells express CTGF in the kidneys. CTGF participates in ECM-associated, profibrotic signaling by binding cell surface receptors (integrins), growth factors (TGFβ, BMPs) and ECM proteins (fibronectin) (Lee, 2012; Yin & Liu, 2019; Rayego-Mateos et al., 2021). Further, both CTGF and PDGF expression is regulated by MRTF/SRF(Sakai et al., 2013; Rozycki et al., 2016; Yamamura et al., 2023), constituting a positive profibrotic feedback loop in the epithelium. IHC quantification confirmed that CTGF and PDGF expression was elevated in the tubules of PKD2 mutant animals compared to those of the control littermates (Figure 8 A, B).

**Figure 8.**
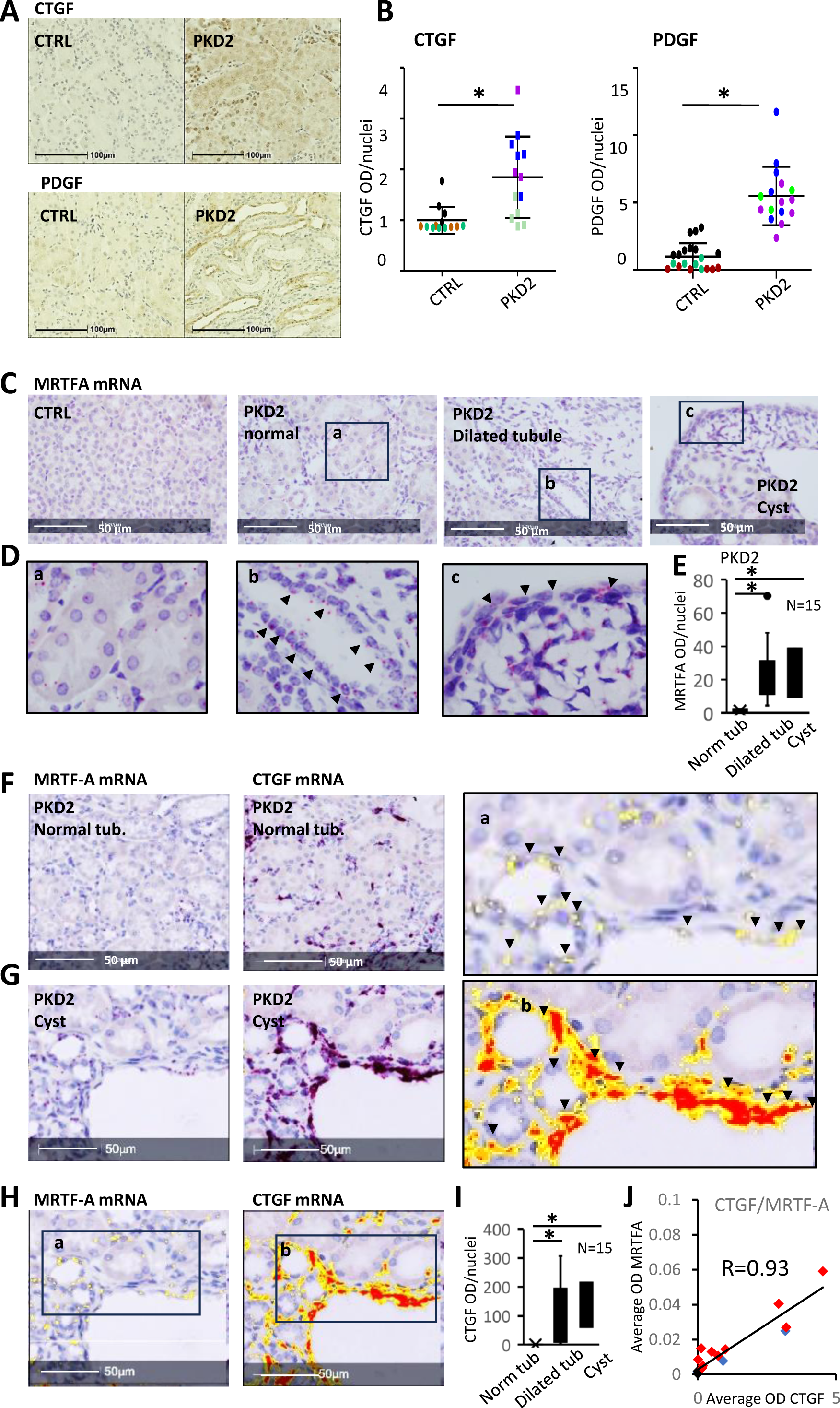
Fibrogenic cytokine expression is increased in PKD, and CTGF expression spatially correlates with MRTF mRNA abundance *in vivo*. (A, B) CTGF and PDGFB expression was detected in PKD2 WS25/- (PKD2) and PKD2 WS25/+ animals (Control, Ctrl) by IHC (n=3). Expression was quantified in 4-10 fields per animal using the Area Quantification package of HALO, normalizing for the number of nuclei. (C-I) RNAScope was used to detect MRTF-A and CTGF mRNA expression in PKD2 WS25/- and PKD2 WS25/+ animals. Images in (D) are enlarged areas of normal and dilated tubules and a cyst. Arrowheads point to MRTF expression signals. HALO Area Quantification software was trained to differentiate the red signal on the hematoxylin-stained background. For easy visualization, positive signals are shown as yellow, orange and red color, corresponding to low, moderate and high signal intensity (H and enlarged areas). RNAscope signal intensity was measured by the Area Quantification module and was normalized to the number of nuclei. (E, I). (J) Correlation of CTGF and MRTF expression is shown in normal- and dilated tubules and cysts of PKD2 WS25/- animals.

We reasoned that if MRTF directly drives CTGF transcription, the corresponding messages should show a spatial correlation; therefore, we compared MRTF-A and CTGF spatial mRNA expression on consecutive sections, using RNAScope. MRTF-A mRNA expression was greatly upregulated in the dilated tubules, the adjacent stroma and the cyst-lining epithelium of PKD2 kidneys, contrasting the minimal expression in normal tubules of the same animals and the control kidneys (Figure 8C-E). Importantly, MRTF-positive epithelial cells ubiquitously expressed CTGF (Figure 8F-I) with a strong positive correlation between tubular MRTF and CTGF expression (Figure 8J). These tubules were typically localized in the vicinity of cysts and stromal areas with strong CTGF expression. Altogether, the mRNA expression of MRTF-A and CTGF spatially overlapped, strengthening the potential role of MRTF in the epithelial initiation of profibrotic signaling (Expanded View Figure 8).

## Discussion

The current study reveals that the RhoA/cytoskeleton/MRTF pathway, a key initiator of epithelial fibrogenesis, is activated in PKD. Thus, MRTF emerges as a significant mediator of PKD-related fibrosis, one of the hallmarks of the disease. MRTF facilitates PEP and the consequent paracrine response, leading to fibroblast-myofibroblast transition in the adjacent stroma. From a broader perspective, PKD is a prominent example of epithelial injury-initiated fibrogenesis, a process that was shown to be sufficient for tubulointerstitial fibrosis (Anorga et al., 2018; Traykova-Brauch et al., 2008; Boutet et al., 2006; Okada et al., 2005; Grgic et al., 2012; Grande et al., 2015).

Nuclear MRTF translocation is the direct consequence for PC loss-induced RhoA activation and cytoskeleton remodeling. We show that knockdown of PC2, similarly to that of PC1 (Anorga et al., 2018; Cai et al., 2018), activates RhoA. Increased RhoA activity can be due to activation of RhoGEFs and/or inhibition of RhoGAPs, and both mechanisms have been proposed in PKD (Streets, Prosseda & Ong, 2020; Cai et al., 2018). How can PC1/PC2 loss affect these processes? One possibility involves heterotrimeric G proteins, particularly G_α12_, which is activated by PC1 loss (Hama & Park, 2016; Parnell et al., 1998; Yuasa et al., 2004; Suzuki, Hajicek & Kozasa, 2009; Suzuki et al., 2003), necessary for PKD pathogenesis (Wu et al., 2016), and can directly stimulate LARG, a GEF implicated in PC1 downregulation-induced RhoA activation (Cai et al., 2018). A non-exclusive alternative entails ITGB1, which is overexpressed in PKD-/- animals, indispensable for cystogenesis and fibrosis (Lee et al., 2015) and can stimulate several GEFs (LARG, GEF-H1) (Guilluy et al., 2011; Shen et al., 2015; Zhang, Reif & Wallace, 2020; Raman et al., 2017).

We show that the downregulation of PC1/2 activates ITGB1 and triggers its partially MRTF-dependent overexpression. While PC loss-induced ITGB1 activation may not be a prerequisite for initial RhoA activation/MRTF induction, sustained ITGB1 activation/expression (promoted by increased cell contractility, stretch of the cyst wall or ECM stiffening (Tanner et al., 1995; Zhang, Reif & Wallace, 2020)) may signify a potent feedforward mechanism. Furthermore, matricellular proteins, such as CTGF (Happé et al., 2011; Gauer et al., 2017) and periostin (Raman et al., 2018, 2017; Wallace et al., 2014), secreted by altered tubules and cysts, can directly activate other integrins, e.g. α_V_β_3_ (Orecchia et al., 2011; Gillan et al., 2002; Gao & Brigstock, 2004), and we show that pan-integrin stimulation is sufficient to provoke strong MRTF nuclear translocation. Taken together, future studies should define critical upstream RhoA-activating mechanisms in PKD, and the contribution of various integrins in the initial and feedforward activation of the RhoA/MRTF pathway.

We show that dysregulated/lost PC signaling induces not only nuclear translocation but also overexpression of MRTF in tubular cells and the cyst-lining epithelium *in vitro* and in PKD1 and 2 mutant animal models. The underlying mechanisms are unknown, but it is worth mentioning that both β-catenin and YAP signaling are upregulated in PKD and other fibrotic diseases, and both can drive MRTF expression (Szeto et al., 2016; Happé et al., 2011; Dwivedi et al., 2020). Conversely, MRTF is a direct inducer of TAZ expression (Speight et al., 2016; Esnault et al., 2014). Indeed, a multilevel crosstalk exist between MRTF and YAP/TAZ wherein these factors can target the same genes and regulate each other’s expression/activity/function (Crider et al., 2011; Fan et al., 2007; Masszi et al., 2010; Speight et al., 2016; Muehlich et al., 2017; Foster, Gualdrini & Treisman, 2017), including MyF differentiation (Dwivedi et al., 2020). Thus, the MRTF-YAP/TAZ synergy constitutes yet another positive feedback loop.

Our targeted gene expression studies and unbiased transcriptome analysis indicated that a substantial cohort of PC1/2 loss-regulated genes are MRTF-sensitive. These encode cytoskeletal components, matricellular proteins/fibrogenic mediators, regulators of cell contractility and integrin signaling, many of which have been implicated in organ scarring. These findings are consistent with previously demonstrated roles of MRTF (reviewed in (Miranda et al., 2021)) and the prominence of SRF, as a key transcriptional hub in PKD(Song et al., 2009b). Moreover, our approaches assigned two novel functions to MRTF in this context. The first regards the expression of genes governing mitochondrial functions/metabolism. This new aspect requires further studies, given that PKD is characterized by robust metabolic changes (reduced OXPHOS, increased glycolysis), and alterations in mitochondrial shape and function (Padovano et al., 2018; Menezes and Germino, 2019; Cassina, Chiaravalli and Boletta, 2020; Podrini, Cassina and Boletta, 2020; Onuchic et al., 2023). The other possible function of MRTF is the *suppression* of inflammatory genes via negative regulation of the relevant transcription factors (e.g. NFκB, as raised before (Wang et al., 2012b; Hayashi et al., 2015)). This may signify a role of MRTF as a switch between the inflammatory and the fibrotic aspects of PKD. In addition, the MRTF-dependent gene sets and GO categories, while overlapping, are not identical for PC1 vs. PC2 loss (Expanded View Figure 4). This highlights PKD-type specific roles of MRTF. While we started to unravel the spatial correlation between MRTF expression and fibrogenic cytokine production, spatial transcriptomics studies should extend these findings in the future.

What is the significance of fibrosis in the overall pathology of PKD? A recent elegant report argues that fibrosis per se may inhibit cyst formation by mechanically restricting cyst growth, but it worsens survival (Zhang et al., 2020). These findings argue that combatting fibrosis is an important therapeutic option in PKD. Our proof-of-principle studies demonstrate that MRTF is activated upon PC loss and in PKD, and suggest that it acts as a significant mediator in the pathobiology of the disease. However, future functional studies should test how genetic or pharmacological interference with MRTF affects various aspects (inflammation, cytogenesis, fibrosis) of the disease, and discern if MRTF is a viable drug target for the treatment of PKD.

## Methods

### Cell lines and reagents

LLC-PK1 (clone 4) a porcine proximal tubule cell line was a gift from Peter Harris (Vanderbilt University School of Medicine, TN) and used as in our previous studies (Masszi et al., 2010). Briefly, cells were cultured in DMEM (11885084, Gibco), supplemented with 10% FBS and penicillin-streptomycin (Life Technologies, Carlsbad, Germany). Transforming growth factor β1 ((TGFβ), R&D, Minneapolis, MN) treatment was carried out for 24 h in serum-free DMEM media at 5 ng/ml concentration.

### siRNA-mediated silencing

siRNA transfections were performed using Lipofectamine RNAiMax (ThermoFisher Scientific), following the manufacturer’s protocol. Cells were transfected at 30-40% confluency. Twenty-four hours post-transfection, cells were serum starved for an additional 24 h, and were assayed at 48 h post-transfection. siRNAs were used at the 100 nM final concentration with exception of siPC1 (150 nM), siRhoA (50 nM) and siMRTF-A (50 nM). Non-related control siRNA (siNR, Silencing Select Negative Control 1, ThermoFisher Scientific) concentration was matched in each experiment.

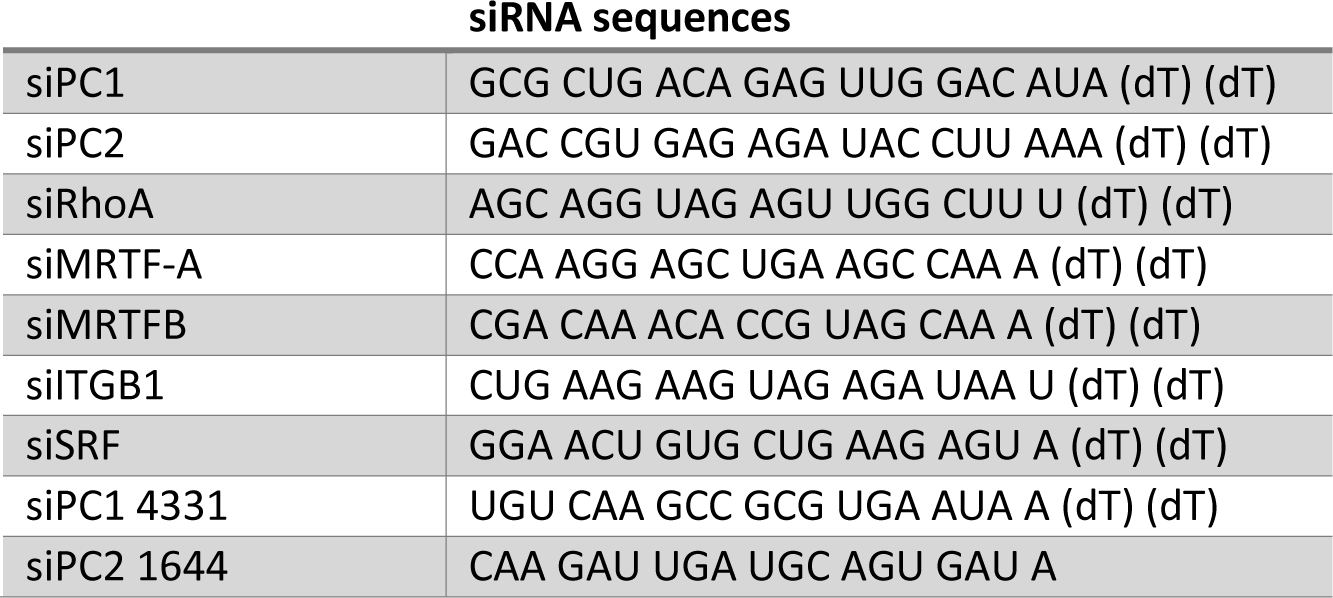

### Real -time quantitative PCR (RT-qPCR)

Total RNA was isolated with RNeasy Mini Kit (Qiagen, Venlo, Netherlands), following the manufacturer’s protocol. 500-1000 ng RNA was reverse transcribed using the iScript Reverse Transcription Supermix (Bio-Rad, Montreal, Canada), and gene expression was quantified using the PowerUP Syber Master Mix (Thermo Fisher Scientific, Waltham, MA, USA) on the QuantStudio 7 Pro system (ThermoFisher Scientific). Relative gene expression was calculated by the ΔΔ Ct method. Melt curve analysis confirmed the amplification of a single product.

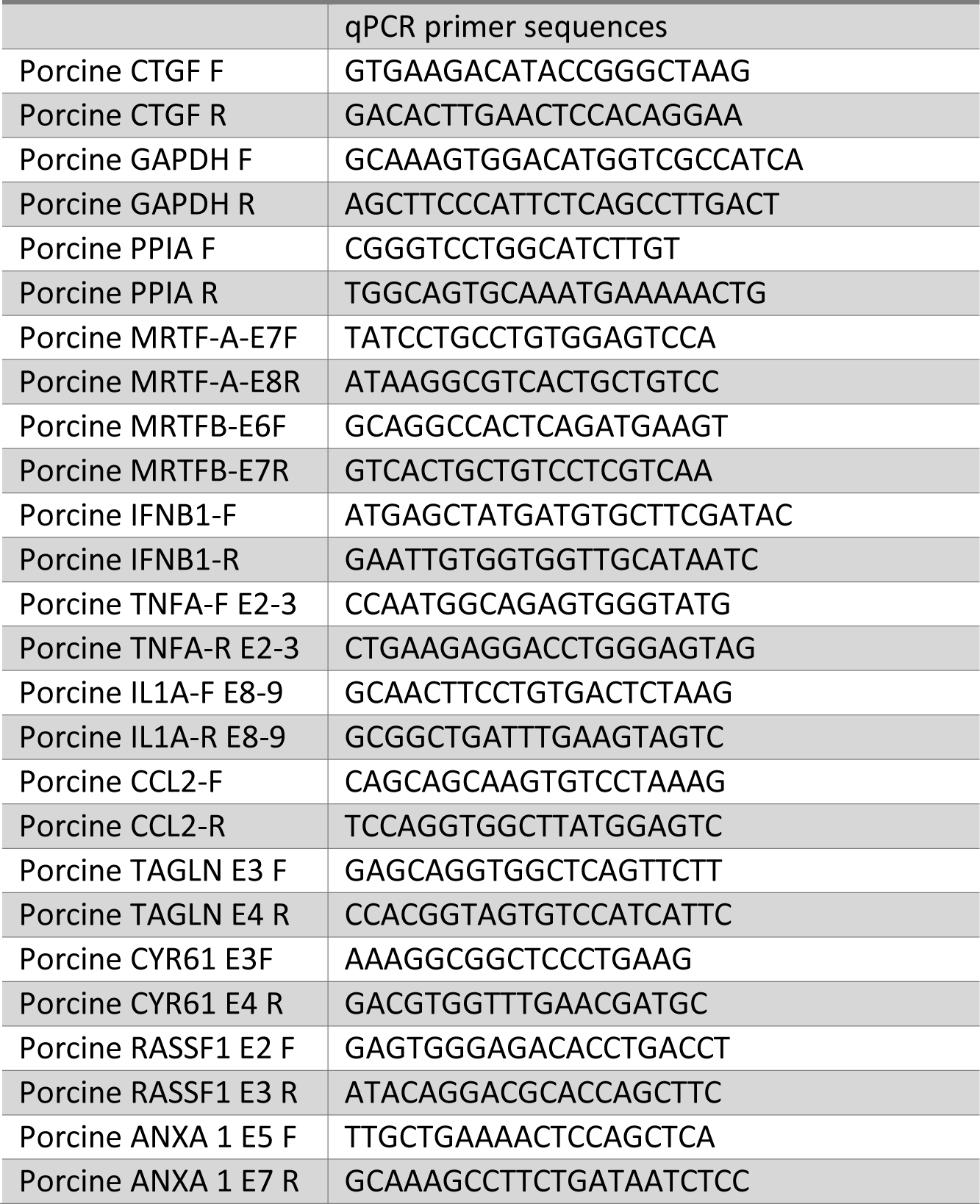

### Western blot analysis

Western blot analysis was performed as described before (Burns & Harris, 1995; Masszi et al., 2010), on nitrocellulose membrane. Blots were visualized by Clarity or Clarity Max ECL Western Blotting Substrate (Bio-Rad Laboratories) and Chemidoc Imaging Systems (Bio-Rad Laboratories). Relative protein expression was calculated by ImageLabs (Bio-Rad). Horseradish peroxidase-conjugated secondary antibodies (1:2000) were from Jackson ImmunoResearch Laboratories. The following primary antibodies were used: Polycystin 1 (1:200 sc-130554, Santa Cruz Biotechnology, TX), Polycystin 2 (1:500, sc-28331, Santa Cruz Biotechnology), MRTF-A (1:1000, s14760 Cell Signaling Technology), and total RhoA (1:1000, 2117S Cell Signaling Technology). GAPDH (sc-47724, Santa Cruz Biotechnology and 6004-1-Ig, Proteintech) or β-actin (A1978, Sigma Aldrich) antibodies were used for normalization.

### Preparation of GST-fusion protein and Rho activation assay

Preparation of GST-RBD (RhoA-binding domain (RBD): amino acids 7-89 of Rhotekin) and the Rho affinity assay were described in (Waheed et al., 2013). The beads were stored frozen in the presence of glycerol. Confluent LLC-PK1 cells were transfected with PC1 or 2 siRNA, and 48 h later lysed with ice-cold assay buffer containing 100 mM NaCl, 50 mM Tris base (pH 7.6), 20 mM NaF, 10 mM MgCl_2_, 1% Triton X-100, 0.5% deoxycholic acid, 0.1% SDS, 1 mM Na_3_VO_4_ and protease inhibitors. The lysates were centrifuged, and aliquots for determining total RhoA were removed. The remaining supernatants were incubated with 20-25 µg of GST-RBD at 4°C for 45 min, followed by extensive washing. Aliquots of total cell lysates and precipitated proteins were analyzed by Western blotting and quantified by densitometry. Precipitated (active) RhoA was normalized using the corresponding total cell lysates.

### Immunofluorescence microscopy and quantification

Cells were grown on glass coverslips for visual expression analysis or on 96-well plates (Corning) for quantitative image analysis (ImagExpress). Cells were transfected as detailed above.

Immunofluorescence staining was performed as described before(Kofler & Kapus, 2021). The following primary antibodies were used: MRTF-A (1:300), active RhoA (1:100, New East Biosciences), active ITGB1 12G10 (1:100 ab150002, Abcam). F-actin was visualized by staining with fluorescently labelled phallodin (Phalloidin iFLUOR 555, Abcam) at 1:10,000 dilution for 1 h. Cells were imaged by either a WaveFX spinning-disk microscopy system (Quorum Technologies) equipped with ORCA-Flash4.0 digital camera or by an Olympus IX81 microscope with the Evolution QEi Monochrome camera, both driven by the MetaMorph software. Nuclear translocation was assessed by ImageXpress, driven by MetaExpress software, as in our previous work (Kofler & Kapus, 2021). Briefly, nuclear staining was measured as mean fluorescence intensity of MRTF-A-specific staining within the DAPI-positive nuclei, and cytoplasmic staining was measured as MRTF-A-positive fluorescence intensity in a preset ring around the nuclei. Nuclear/cystoplasmic ratio of MRTF-A was arranged in bins incremented by 0.02 (x axis). Active integrin clusters were counted by the Metamorph software using the Manually Count Objects option. Counts were normalized to cell number.

### Next Generation Sequencing transcriptome analysis

mRNA libraries were prepared using the NEBNext® Poly(A) mRNA Magnetic Isolation Module, NEBNext® Ultra™ II Directional RNA Library Prep with Sample Purification Beads and NEBNext Multiplex Oligos for Illumina (96). Sequencing was carried out on the NovaSeq SP flowcell SR200 or PE100. Data analysis is detailed in Expanded View Methods.

### LLC-PK1 and fibroblast communication, collagen substrate wrinkling quantification

Wild type subcutaneous fibroblasts (WT SCF) were isolated from C57BL/6 mice and cultured on soft substrates with a Young’s elastic modulus of 0.2 kPa. Substrates were generated as described(Goffin et al., 2006) Subsequently, substrates were oxygenized and coated with gelatin (2ug/cm^2^ diluted in PBS)(Wipff et al., 2009) 2.000 SCF/cm^2^ were seeded and maintained in Dulbecco’s modified Eagle’s medium (DMEM) supplemented with 10% fetal bovine serum (FBS) and 1% penicillin/streptomycin for 6 h prior synchronization with serum-free DMEM overnight. Conditioned media (CM) was obtained from LLC-PK1 PK1 renal epithelial cells. Epithelial cells were transfected with the indicated siRNAs. Twenty-four h post-transfection, the media was removed, cells were washed three times and fresh serum-free media was added. After 24 h (48 h post-transfection), CM was collected. Fibroblasts were stimulated with the CM for 48 h. Subsequently, phase contrast images were taken of the fibroblasts by using Zeiss Axio Observer Microscope. Cell contractile function, related to % area covered by wrinkles, was analyzed by FIJI ImageJ software and the values were normalized to cell number.

### Animal tissues

Animal housing and experimental protocols were carried out in accordance with the Research Ethics Board of University Health Network, Toronto. Paraffin-embedded blocks of the kidneys of 12 months old PKD1 RC/RC (n=3) and control littermates (WT, n=3), as well as 3 months old PKD2 WS25/- (n=6) and control littermates (PKD2 WS25/+, n=3) (Hopp et al., 2012; Cai et al., 2018; Wu et al., 1998) were processed at the Keenan Research Centre for Biomedical Sciences, St Michael’s Hospital.

### Patient specimens

Nephrectomy specimens were collected from PKD and RCC patients, following their informed consent. All protocols were approved by the Research Ethics Board of the Keenan Research Centre for Biomedical Sciences, St Michael’s Hospital (REB #20-295).

### Immunohistochemistry and quantification

Paraffin-embedded sections were stained as. Antigen retrieval was performed by boiling the section for 5 min in TE buffer. Staining was viewed by Olympus BX50 microscope, using the Cellsense software. Stained sections were scanned with Axio Scan Z1 driven by the ZEN software and analyzed by HALO v2.3. For quantifying CTGF and PDGF-B, 3 animals in each group were used, and DAB staining was quantified by HALO in 5-10 large, randomly selected fields. DAB OD was normalized to the number of nuclei in each area. For quantitation of MRTF-A staining, thresholds were set to recognize hematoxylin stain (nuclei) and DAB (MRTF-A), and average DAB staining intensity was determined for each nucleus. The total scale of DAB OD was divided into 256 evenly distributed intensity bins (X axis) and frequency for each intensity bin was calculated in Excel. Six PKD2 WS25/- animals and 6 controls (PKD2 WS25/+ littermates) were analyzed, however 1 PKD2 WS25/- animal was removed from the quantification, because it did not develop renal cysts. Tubules with over 1.5x diameter of normal tubules were classified as dilated. Microcysts had flattened epithelial lining and, in general, lacked brush border.

### RNAScope

Three months old PKD2 WS25/- (n=3) and control PKD2 WS25/+ littermates (n=3) were analyzed for MRTF-A and CTGF mRNA expression, using 3 μm thin sections of paraffin embedded kidneys. Staining was carried out by closely following the manufacturer’s protocol; 10% Gill’s hematoxylin was used to stain nuclei. Stained sections were scanned by Axio Scan Z1 and were analyzed using the HALO software v2.3. Renal tubules with normal and dilated morphology (n=15), and cysts (n=5) were selected for analysis. Thresholds were set for hematoxylin, MRTF-A and CTGF staining. OD was normalized to the number of nuclei.

### Statistical analysis

Immunofluorescence and Western blot images show representative results of minimum three similar experiments or the number of experiments indicated (n). Graphs show the means ± standard error of the mean. Statistical significance was determined by Student’s *t*-test or one-way analysis of variance (Tukey posthoc testing), using Prism software. Violin plots were generated with the statskingdom.com/violin-plot-maker.html website, including the median and excluding outliers (Tukey) options. Statistical significance was calculated using one-sample or two-tailed t-test, as appropriate. *P*<0.05 was accepted as significant. Unless indicated otherwise, *, ** and *** correspond to *p*<0.05, <0.01 and <0.001, respectively.

## Acknowledgement

We are indebted to Dr. McCulloch for providing the 12G10 antibody, and Drs. Di Ciano-Oliveira and Pamela Plant for advice and help with image analysis and mRNA library quantification.

This work was supported by grants from the Canadian Institutes of Health Research (CIHR, PJT 178152, PJT 162360) to AK and (CIHR, PJT 180482) to KS, and the Kidney Foundation of Canada to AK.

AK is a Thor Eaton Professor of Fibrosis Research, jointly appointed by the University of Toronto and the St. Michael’s Hospital.

## Expanded View Figure Legends

**Expanded View Figure 1.**
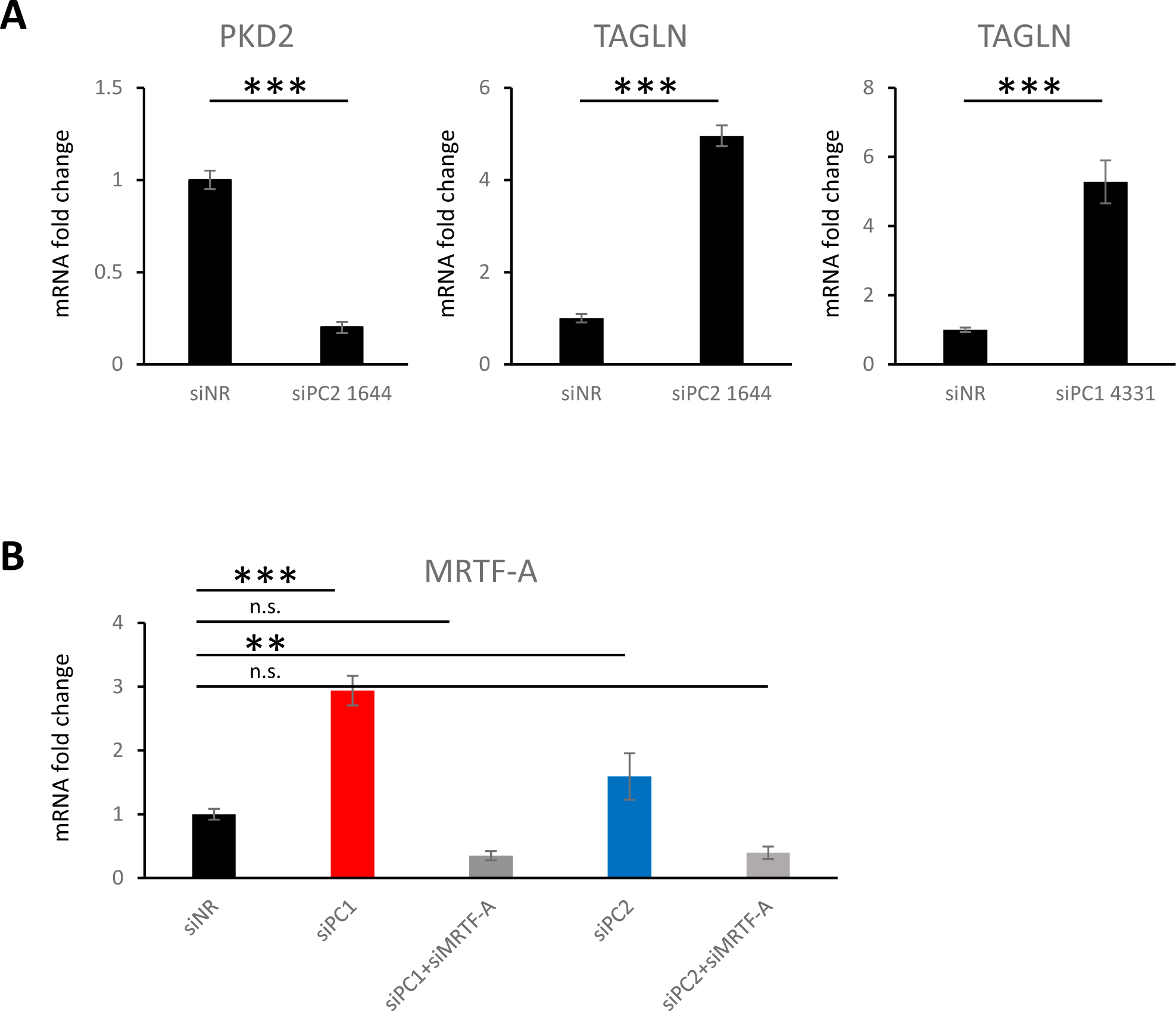
Experimental setup. (A) siPC1-1433 and siPC2-1644 are a second set of PC1 and PC2-targeting siRNAs (respectively). Transfection of either siPC1-1433 (150 nM) or PC2-1644 (100 nM) enhanced TAGLN expression. qPCR data is shown. (B) MRTF-A mRNA was quantified by RT-qPCR. siPC1 or siPC2 knockdown did not alter the efficiency of MRTF-A silencing (n=3, n.s. non-significant, * p<0.05, ** p<0.01).

**Expanded View Figure 2.**
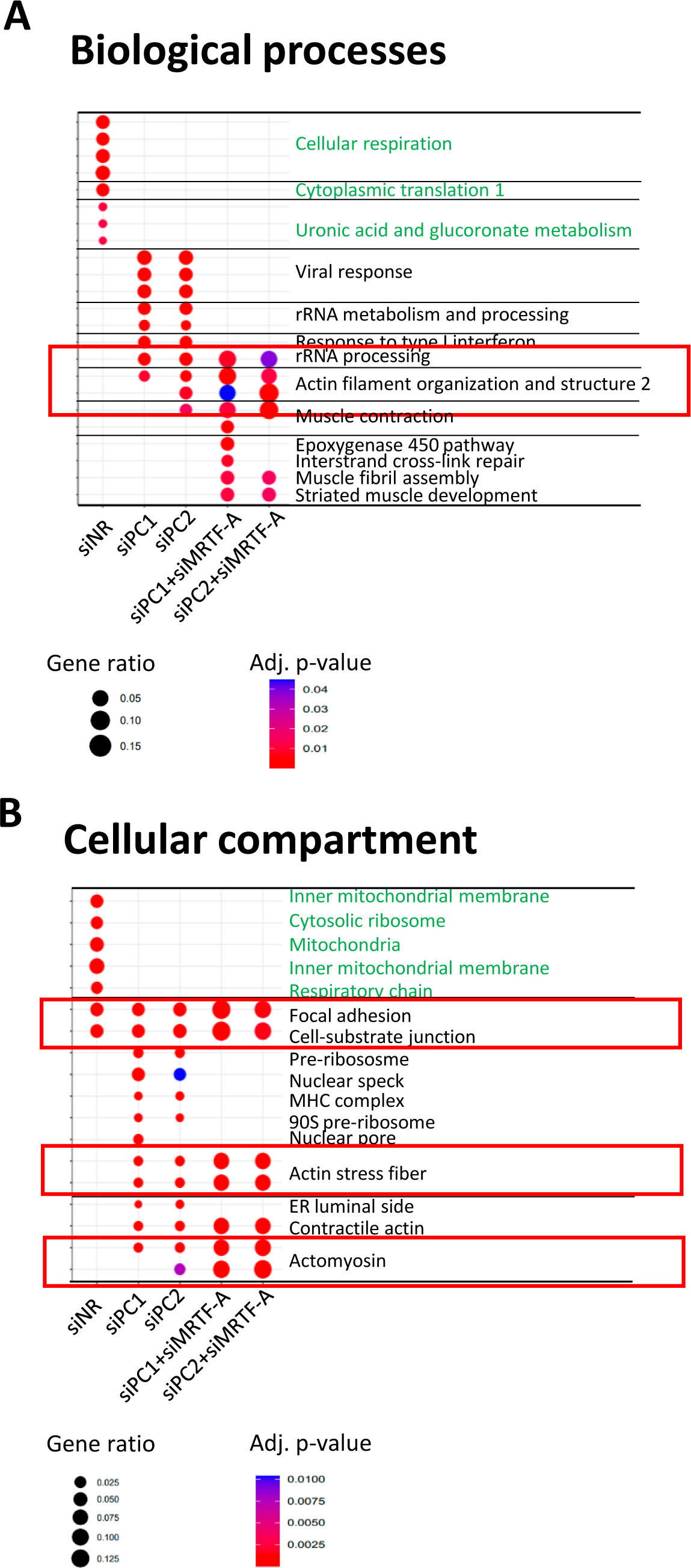
PKD-linked cytoskeletal changes are regulated by MRTF. Differentially expressed genes were determined as detailed in Expanded View Methods and were subjected to enrichment analysis of Biological Processes (A) and Cellular Compartment (B), using ClusterProfiler in R. Categories that were depleted in siPC1 and siPC2 conditions (compared to siNR) are shown in green. Further, selected top categories that were enriched in siPC1 or siPC2 conditions (compared to siNR) are listed in the siPC1 and siPC2 columns. Significant decrease compared to these in siMRTF-treated samples are shown in the next two columns. Note that biological processes that overlap between PC knockdown conditions and the siPC+siMRTF-A conditions were related to cytoskeleton organization and cell contractility (red rectangles).

**Expanded View Figure 3.**
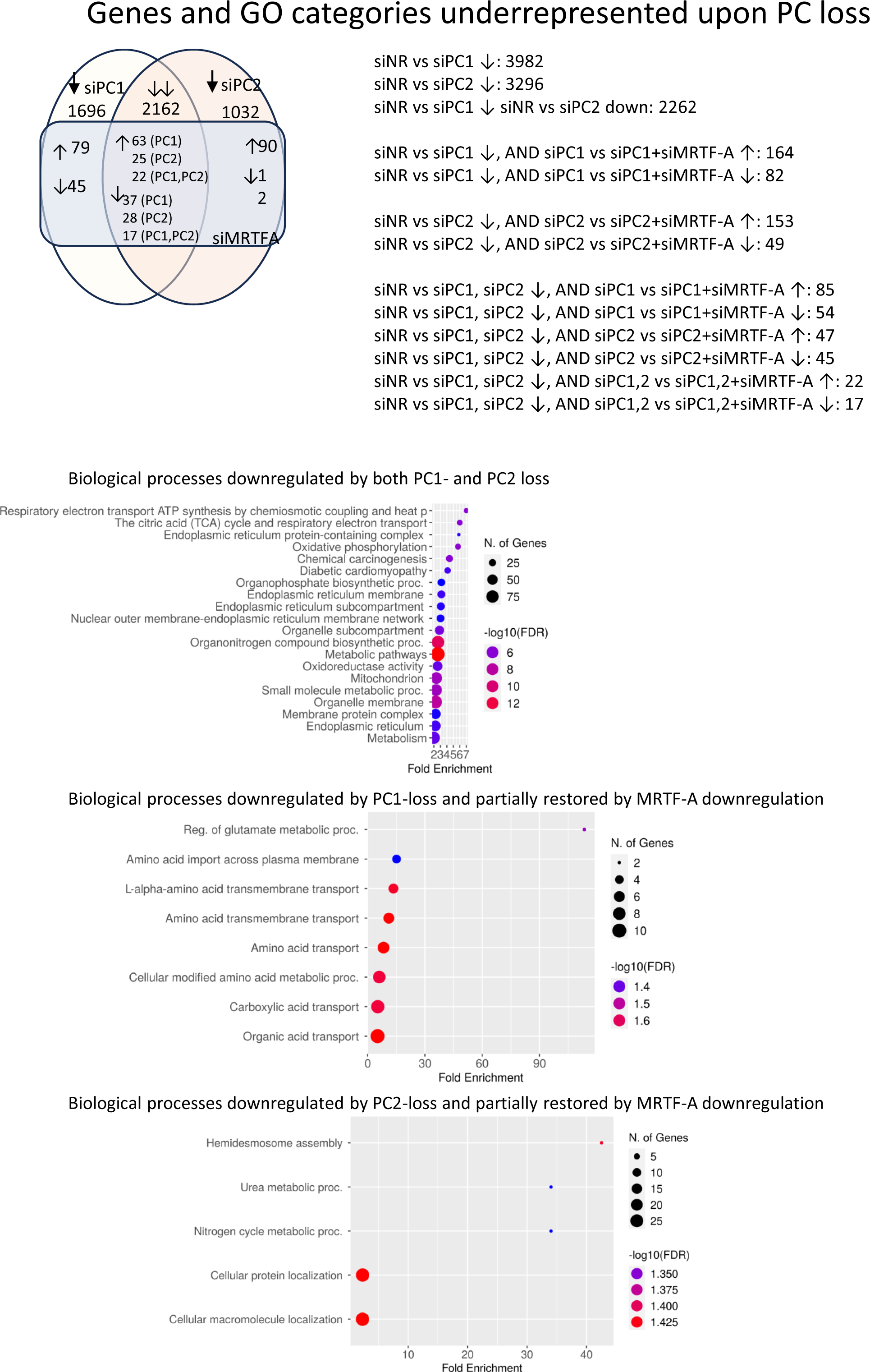
PC1 and PC2 loss-specific underrepresented biological processes and their MRTF-dependence. Top panel: Epithelial cells knocked down for PC1 and PC2 transcriptionally underrepresented several metabolic pathways. Lower panels: Out of these, MRTF-A silencing partially mitigates amino acid transport across plasma membrane (siPC1).

**Expanded View Figure 4.**
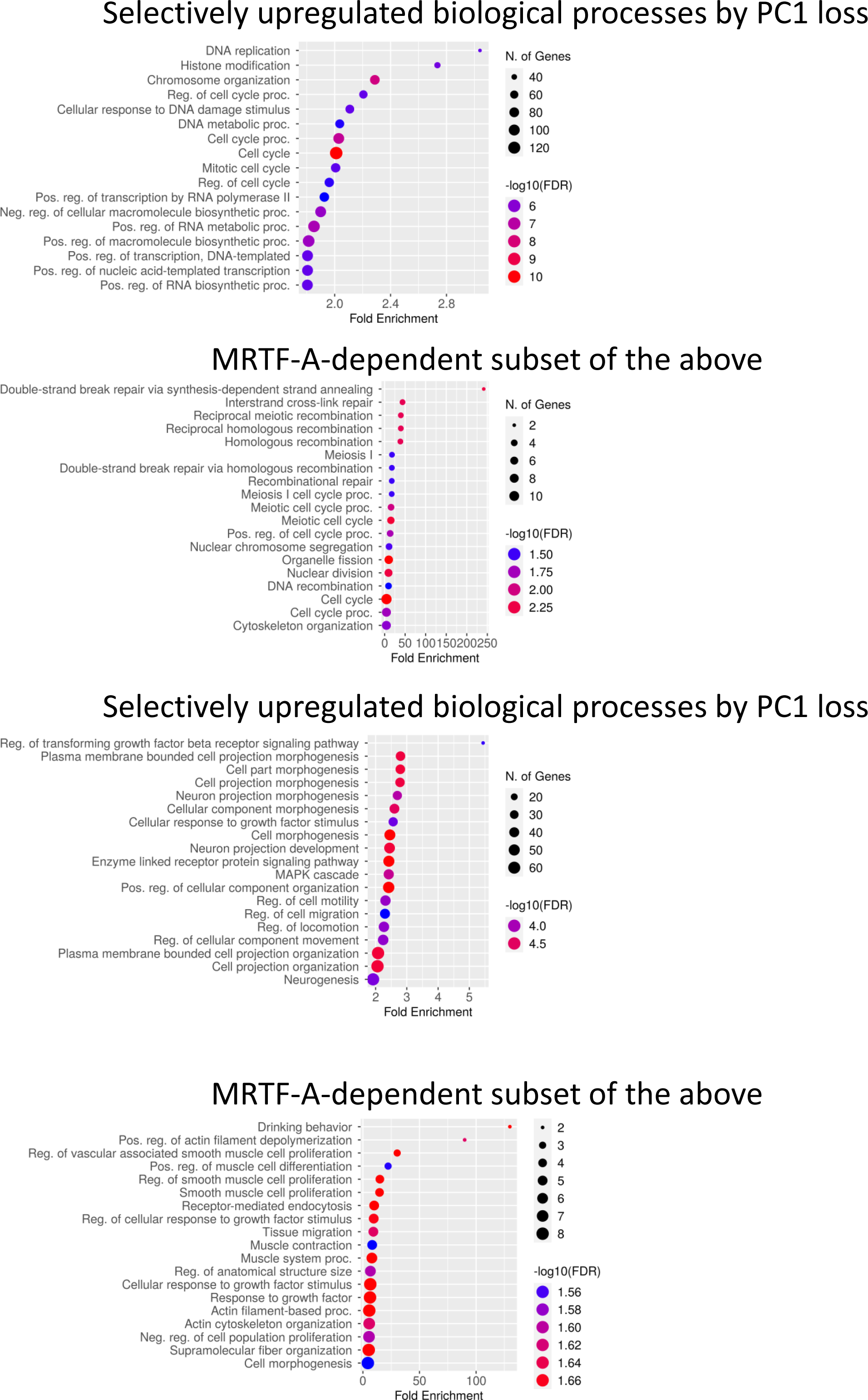
PC1 and PC2 loss-specific biological processes. DEG searched for transcripts that were upregulated upon PC1 loss but were unaffected by PC2 loss, or vice versa. For PC1 loss-related biological processes centered around DNA replication and transcription (top panel). Within this transcript set, MRTF-dependent genes were grouped in DNA recombination-related repair. Interestingly, PC2-specific upregulated categories remained to be linked to cell projection/cytoskeletal reorganization/cell motility. The MRTF-dependent subset is predicted to regulate the actin cytoskeleton, cell contractility and muscle development.

**Expanded View Figure 5.**
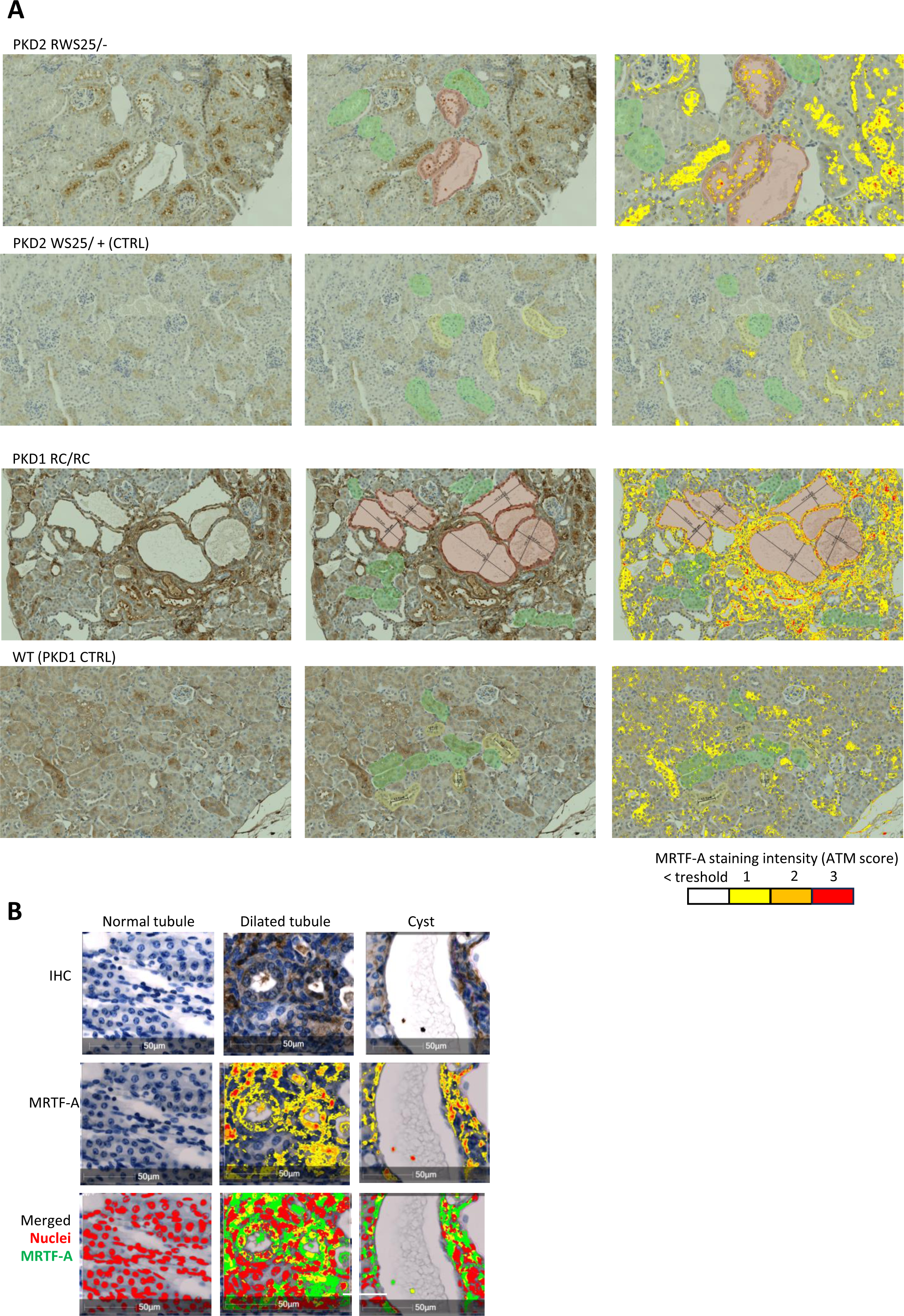
Expression analysis methods. (A) ATM within the Area Quantification Module measured MRTF-A/DAB signal and colored each pixel according to its DAB optical density. Yellow to red color gradient corresponds to increasing MRTF-A expression, as indicated. Green and yellow masks show selected normal tubules without or with lumen, respectively. Red masks depict visibly enlarged tubules; lumen measurements are given in μm. (B) Automated Area Quantification Module was trained to recognize the hematoxylin-stained nuclei and to depict MRTF-A expression, as in (A) (middle row panels). The bottom panels show a superimposed image of the nuclear mask and the MRTF-A-specific stain. Here red indicates nuclei, green indicates MRTF-A and yellow depicts nuclear MRTF-A.

**Expanded View Figure 6.**
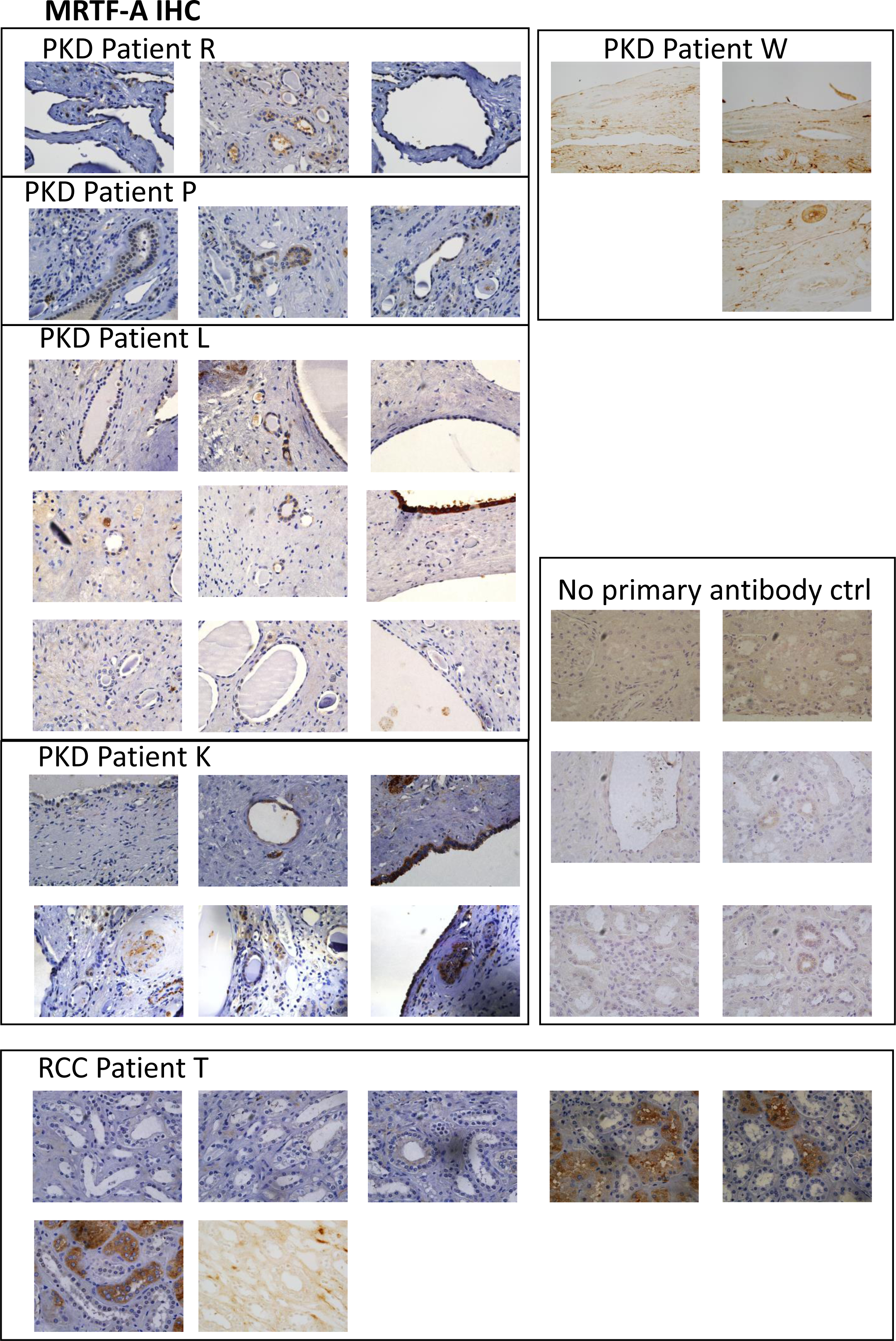
ADPKD patients’ cystic epithelium shows enhanced MRTF-A expression. Five ADPKD patients (patient R, P, L, K, W) and control (normal kidney tissue from a clear cell renal cell carcinoma patient) specimens were assessed for MRTF-A expression by IHC. Nuclear expression of MRTF-A was observed in all ADPKD patients, albeit the extent varied. The cystic epithelial wall showed enhanced MRTF expression compared to the normal tubules within the same patient’s specimens and to the normal tubules of the RCC patient.

**Expanded View Figure 7.**
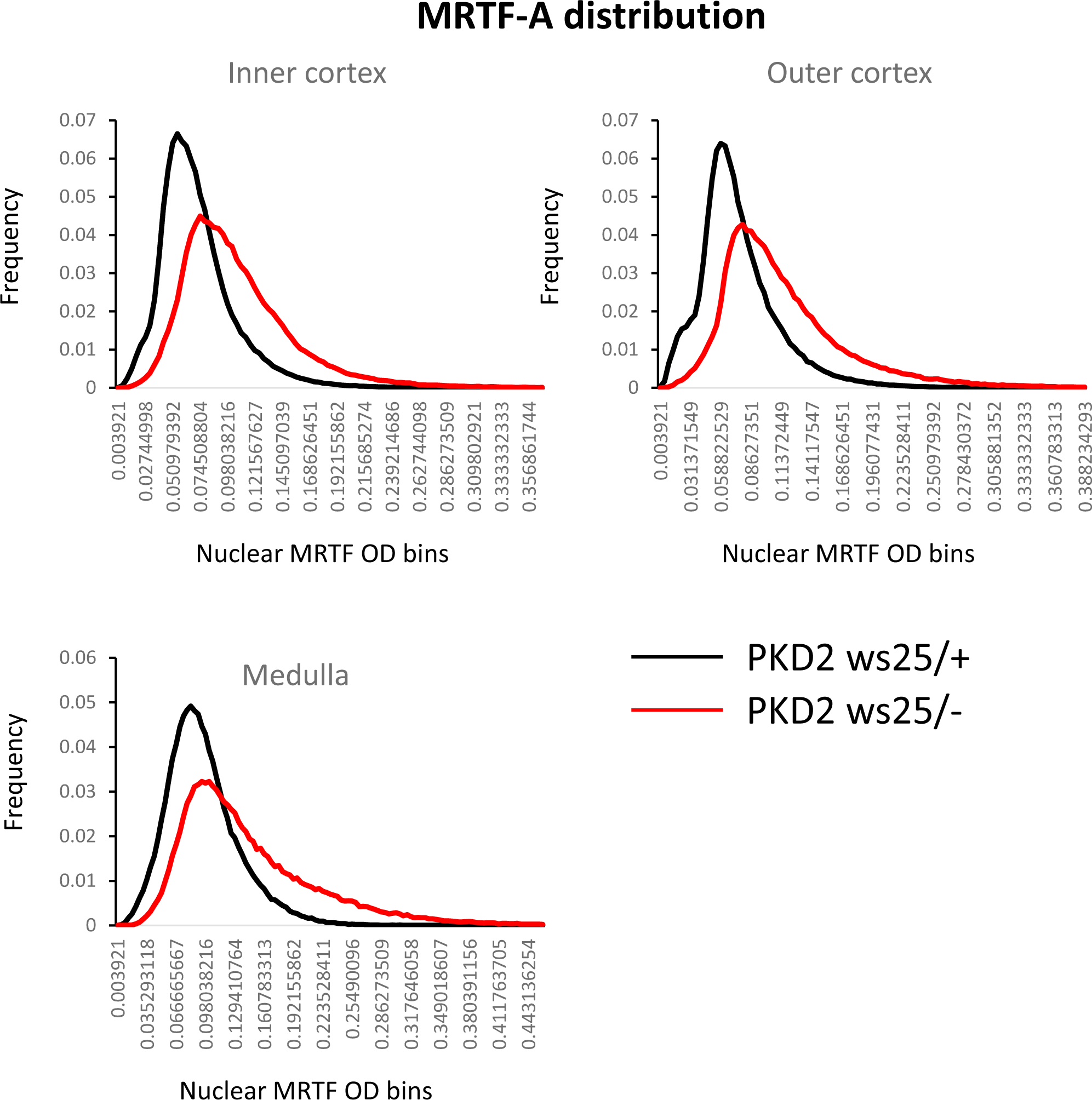
Distribution curves show the frequency of nuclei with increasing MRTF-A expression. The inner cortex, outer cortex and medulla was assessed separately in PKD2 WS25/- (3 months old, n=5) and control PKD2 WS25/+ (3 months old, n=6) animals.

**Expanded View Figure 8.**
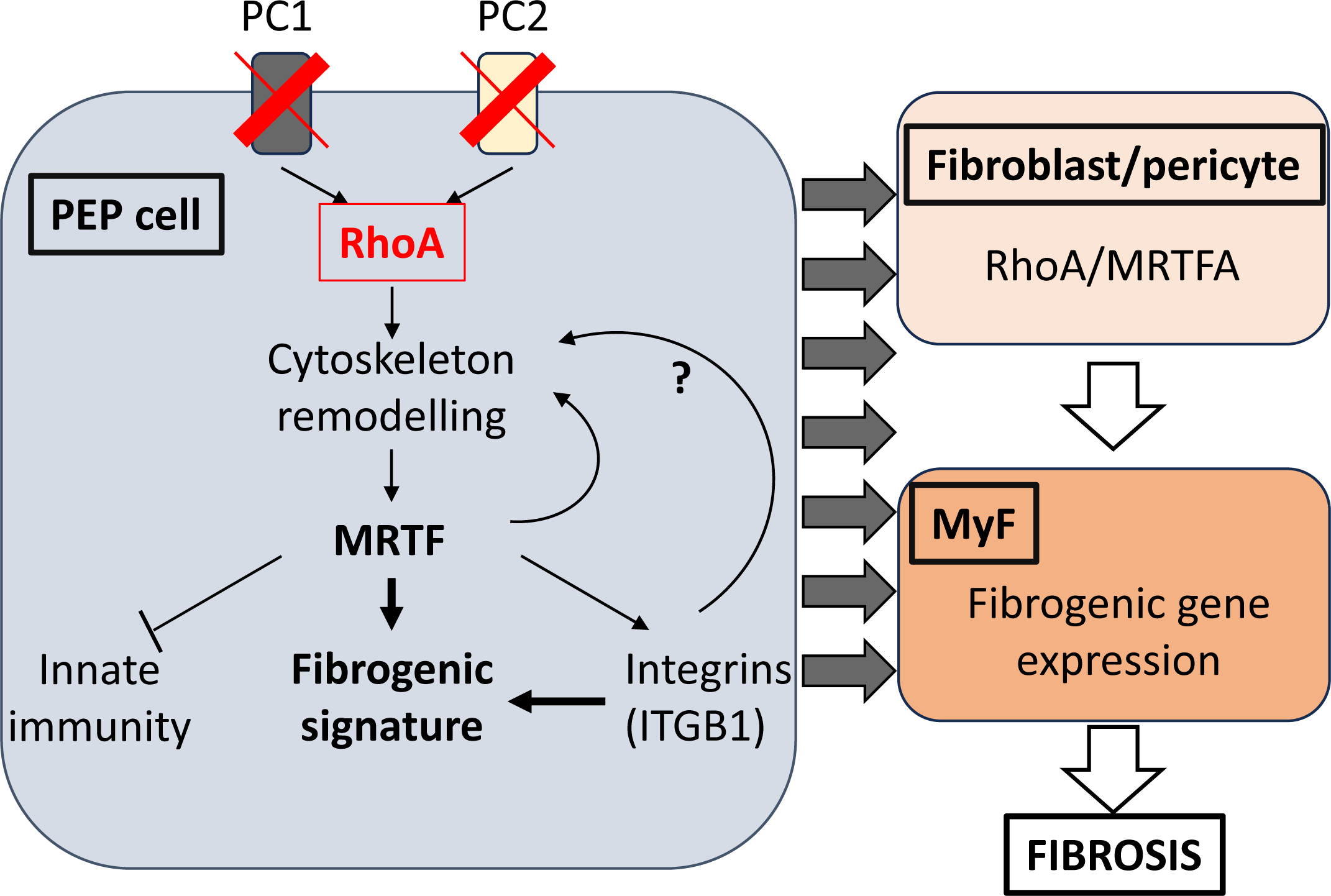
Potential framework of MRTF-related fibrogenic processes in PKD. In resting epithelial cells, the transcriptional co-activator MRTF is bound to globular actin monomers that mask its nuclear localization signal and promote its efflux from the nucleus. Thus, MRTF is retained in the cytoplasm. Loss of PCs result in the activation of RhoA, a master regulator of the cytoskeleton. The ensuing actin polymerization leads to the release of G-actin from MRTF, thereby facilitating its nuclear translocation and accumulation. MRTF binds to SRF and initiates the transcription of CArG-box-dependent genes that regulate the expression of cytoskeletal and matricelllular genes, promoting fibrosis. One of these, ITGB1 has been shown to play a key role in the pathogenesis of PKD. Moreover, ITGB1 facilitates MRTF’s nuclear translocation, creating and MRTF-ITGB1 positive feedback loop, which may contribute to sensitizing epithelial cells for injury. On the other hand, MRTF seems to inhibit TNFα-related proinflammatory signals in the epithelium; however, MRTF reportedly enhances inflammatory signal in macrophages. The MRTF-SRF-mediated transcriptional reprogramming leads to a ‘profibrotic epithelial phonotype’ (PEP), a partial EMT state that is specialized for autocrine and paracrine profibrotic signaling. PEP paracrine signals induce the transition of stromal elements (pericytes and fibroblasts) to myofibroblasts (MyFs), a contractile cell state that is critical for wound healing. However, excessive accumulation and activation of MyFs leads to robust extracellular matrix deposition and fibrosis.

## Expanded View Methods

A series of preprocessing steps were used to ensure RNA-seq data quality. FastQC v0.11.8 for quality control assessment and Trimgalore v0.6.3_dev (https://www.bioinformatics.babraham.ac.uk/projects/trim_galore/) to eliminate low-quality reads. For quantification of expressed transcripts, a quasi-mapping Salmon 0.14.1 was used mapping to the pig reference transcriptome (Ensembl: Sus_scrofa.Sscrofa11.1.cds.all 2023-04-22)^8^. Transcript-level abundances were aggregated to the gene level using "tximport 1.12.3"^9^ and genes with limited read counts were filtered out, requiring genes to have a count in at least 2 samples equal to the smallest group size. Principal components analysis (PCA) was used to assess the overall similarity between samples.

Obtained mapped reads were analyzed with Weighted Gene Co-expression Network Analysis (WGCNA) and Differential Gene Expression (DGE) analyses. For the WGCNA, soft threshold powers were assessed, and a power of 7 was chosen to create a scale-free topology model using default parameters. This model generated a topological overlap and adjacency matrix, facilitating the identification of gene modules through hierarchical clustering. Gene modules comprises of closely interconnected genes with shared co-expression patterns. The module eigengene (ME) within a module serves as the first principal component, providing a representation of the overall expression level for that specific module^10^.

DGE analysis was performed using DESeq2 1.24.0^11^, with significant genes identified based on a false discovery rate (FDR) threshold of 5% and an absolute log2 fold change threshold of 0.6. The Hochberg (FDR) method was used to assign p-values for stepwise/sequential analysis. An alternative approach involved aligning expressed genes to the Sus scrofa reference genome (Sus scrofa 11.1, http://asia.ensembl.org/Sus_scrofa/Info/Index) using the HISAT2 aligner^12^. Transcript abundance was then calculated using an R Bioconductor^13^ in conjunction with the Rsubread package^14^. The mapped reads were also investigated with DEXSeq bioconductor package to identify instances where exons show variations in exon usage^15^.

Gene Set Enrichment Analysis (GSEA) (Software: http://www.broad.mit.edu/gsea/)^16,17^ with the Molecular Signature Database (Human MSigDB v2023.1. Hs updated March 2023) was performed with default parameters to identify potential gene pathways^11^. Candidate genes from the different analyses were subjected for enrichment pathway databases, including Hallmark Gene Sets, Positional Gene Sets, GO Gene Sets, KEGG, and All Transcription Factor Targets, using ShinyGo^18,19^.

## References

Anorga, S., Overstreet, J.M., Falke, L.L., Tang, J., Goldschmeding, R.G., Higgins, P.J. & Samarakoon, R. (2018) Deregulation of Hippo-TAZ pathway during renal injury confers a fibrotic maladaptive phenotype. FASEB journal : official publication of the Federation of American Societies for Experimental Biology. 32 (5), 2644–2657. doi:10.1096/fj.201700722R.

Bergmann, C., Guay-Woodford, L.M., Harris, P.C., Horie, S., Peters, D.J.M. & Torres, V.E. (2018) Polycystic kidney disease. Nature reviews. Disease primers. 4 (1), 50. doi:10.1038/s41572-018-0047-y.

Bialik, J.F., Ding, M., Speight, P., Dan, Q., Miranda, M.Z., Di Ciano-Oliveira, C., Kofler, M.M., Rotstein, O.D., Pedersen, S.F., Szászi, K. & Kapus, A. (2019) Profibrotic epithelial phenotype: a central role for MRTF and TAZ. Scientific reports. 9 (1), 4323. doi:10.1038/s41598-019-40764-7.

Boutet, A., De Frutos, C.A., Maxwell, P.H., Mayol, M.J., Romero, J. & Nieto, M.A. (2006) Snail activation disrupts tissue homeostasis and induces fibrosis in the adult kidney. The EMBO journal. 25 (23), 5603– 5613. doi:10.1038/sj.emboj.7601421.

Brandt, D.T., Baarlink, C., Kitzing, T.M., Kremmer, E., Ivaska, J., Nollau, P. & Grosse, R. (2009a) SCAI acts as a suppressor of cancer cell invasion through the transcriptional control of beta1-integrin. Nature cell biology. 11 (5), 557–568. doi:10.1038/ncb1862.

Brandt, D.T., Xu, J., Steinbeisser, H. & Grosse, R. (2009b) Regulation of myocardin-related transcriptional coactivators through cofactor interactions in differentiation and cancer. *Cell cycle (Georgetown*, Tex.). 8 (16), 2523–2527. doi:10.4161/cc.8.16.9398.

Burns, K.D. & Harris, R.C. (1995) Signaling and growth responses of LLC-PK1/Cl4 cells transfected with the rabbit AT1 ANG II receptor. The American journal of physiology. 268 (4 Pt 1), C925–35. doi:10.1152/ajpcell.1995.268.4.C925.

Cai, J., Song, X., Wang, W., Watnick, T., Pei, Y., Qian, F. & Pan, D. (2018) A RhoA-YAP-c-Myc signaling axis promotes the development of polycystic kidney disease. Genes & development. 32 (11–12), 781–793. doi:10.1101/gad.315127.118.

Caplan, M.J. (2022) AMPK and Polycystic Kidney Disease Drug Development: An Interesting Off-Target Target. Frontiers in medicine. 9, 753418. doi:10.3389/fmed.2022.753418.

Cassina, L., Chiaravalli, M. & Boletta, A. (2020) Increased mitochondrial fragmentation in polycystic kidney disease acts as a modifier of disease progression. FASEB journal : official publication of the Federation of American Societies for Experimental Biology. 34 (5), 6493–6507. doi:10.1096/fj.201901739RR.

Chapin, H.C. & Caplan, M.J. (2010) The cell biology of polycystic kidney disease. The Journal of cell biology. 191 (4), 701–710. doi:10.1083/jcb.201006173.

Crider, B.J., Risinger, G.M., Haaksma, C.J., Howard, E.W. & Tomasek, J.J. (2011) Myocardin-related transcription factors A and B are key regulators of TGF-β1-induced fibroblast to myofibroblast differentiation. The Journal of investigative dermatology. 131 (12), 2378–2385. doi:10.1038/jid.2011.219.

Doke, T., Abedini, A., Aldridge, D.L., Yang, Y.-W., Park, J., et al. (2022) Single-cell analysis identifies the interaction of altered renal tubules with basophils orchestrating kidney fibrosis. Nature immunology. 23 (6), 947–959. doi:10.1038/s41590-022-01200-7.

Douguet, D., Patel, A. & Honoré, E. (2019) Structure and function of polycystins: insights into polycystic kidney disease. Nature reviews. Nephrology. 15 (7), 412–422. doi:10.1038/s41581-019-0143-6.

Dwivedi, N., Tao, S., Jamadar, A., Sinha, S., Howard, C., Wallace, D.P., Fields, T.A., Leask, A., Calvet, J.P. & Rao, R. (2020) Epithelial Vasopressin Type-2 Receptors Regulate Myofibroblasts by a YAP-CCN2-Dependent Mechanism in Polycystic Kidney Disease. Journal of the American Society of Nephrology : JASN. 31 (8), 1697–1710. doi:10.1681/ASN.2020020190.

Esnault, C., Stewart, A., Gualdrini, F., East, P., Horswell, S., Matthews, N. & Treisman, R. (2014) Rho-actin signaling to the MRTF coactivators dominates the immediate transcriptional response to serum in fibroblasts. Genes & development. 28 (9), 943–958. doi:10.1101/gad.239327.114.

Fan, L., Sebe, A., Péterfi, Z., Masszi, A., Thirone, A.C.P., Rotstein, O.D., Nakano, H., McCulloch, C.A., Szászi, K., Mucsi, I. & Kapus, A. (2007) Cell contact-dependent regulation of epithelial-myofibroblast transition via the rho-rho kinase-phospho-myosin pathway. Molecular biology of the cell. 18 (3), 1083– 1097. doi:10.1091/mbc.e06-07-0602.

Ferreira, F.M., Watanabe, E.H. & Onuchic, L.F. (2015) Polycystins and Molecular Basis of Autosomal Dominant Polycystic Kidney Disease.

Fintha, A., Gasparics, Á., Fang, L., Erdei, Z., Hamar, P., Mózes, M.M., Kökény, G., Rosivall, L. & Sebe, A. (2013) Characterization and role of SCAI during renal fibrosis and epithelial-to-mesenchymal transition. The American journal of pathology. 182 (2), 388–400. doi:10.1016/j.ajpath.2012.10.009.

Foster, C.T., Gualdrini, F. & Treisman, R. (2017) Mutual dependence of the MRTF-SRF and YAP-TEAD pathways in cancer-associated fibroblasts is indirect and mediated by cytoskeletal dynamics. Genes & development. 31 (23–24), 2361–2375. doi:10.1101/gad.304501.117.

Fragiadaki, M., Macleod, F.M. & Ong, A.C.M. (2020) The Controversial Role of Fibrosis in Autosomal Dominant Polycystic Kidney Disease. International journal of molecular sciences. 21 (23). doi:10.3390/ijms21238936.

Gao, R. & Brigstock, D.R. (2004) Connective tissue growth factor (CCN2) induces adhesion of rat activated hepatic stellate cells by binding of its C-terminal domain to integrin alpha(v)beta(3) and heparan sulfate proteoglycan. The Journal of biological chemistry. 279 (10), 8848–8855. doi:10.1074/jbc.M313204200.

Gauer, S., Holzmann, Y., Kränzlin, B., Hoffmann, S.C., Gretz, N., Hauser, I.A., Goppelt-Struebe, M., Geiger, H. & Obermüller, N. (2017) CTGF Is Expressed During Cystic Remodeling in the PKD/Mhm (cy/+) Rat Model for Autosomal-Dominant Polycystic Kidney Disease (ADPKD). The journal of histochemistry and cytochemistry : official journal of the Histochemistry Society. 65 (12), 743–755. doi:10.1369/0022155417735513.

Gillan, L., Matei, D., Fishman, D.A., Gerbin, C.S., Karlan, B.Y. & Chang, D.D. (2002) Periostin secreted by epithelial ovarian carcinoma is a ligand for alpha(V)beta(3) and alpha(V)beta(5) integrins and promotes cell motility. Cancer research. 62 (18), 5358–5364.

Goffin, J.M., Pittet, P., Csucs, G., Lussi, J.W., Meister, J.-J. & Hinz, B. (2006) Focal adhesion size controls tension-dependent recruitment of alpha-smooth muscle actin to stress fibers. The Journal of cell biology. 172 (2), 259–268. doi:10.1083/jcb.200506179.

Gopalakrishnan, J., Feistel, K., Friedrich, B.M., Grapin-Botton, A., Jurisch-Yaksi, N., Mass, E., Mick, D.U., Müller, R.-U., May-Simera, H., Schermer, B., Schmidts, M., Walentek, P. & Wachten, D. (2023) Emerging principles of primary cilia dynamics in controlling tissue organization and function. The EMBO journal. 42 (21), e113891. doi:10.15252/embj.2023113891.

Grande, M.T., Sánchez-Laorden, B., López-Blau, C., De Frutos, C.A., Boutet, A., Arévalo, M., Rowe, R.G., Weiss, S.J., López-Novoa, J.M. & Nieto, M.A. (2015) Snail1-induced partial epithelial-to-mesenchymal transition drives renal fibrosis in mice and can be targeted to reverse established disease. Nature medicine. 21 (9), 989–997. doi:10.1038/nm.3901.

Grgic, I., Campanholle, G., Bijol, V., Wang, C., Sabbisetti, V.S., Ichimura, T., Humphreys, B.D. & Bonventre, J. V (2012) Targeted proximal tubule injury triggers interstitial fibrosis and glomerulosclerosis. Kidney international. 82 (2), 172–183. doi:10.1038/ki.2012.20.

Gualdrini, F., Esnault, C., Horswell, S., Stewart, A., Matthews, N. & Treisman, R. (2016) SRF Co-factors Control the Balance between Cell Proliferation and Contractility. Molecular cell. 64 (6), 1048–1061. doi:10.1016/j.molcel.2016.10.016.

Guilluy, C., Swaminathan, V., Garcia-Mata, R., O’Brien, E.T., Superfine, R. & Burridge, K. (2011) The Rho GEFs LARG and GEF-H1 regulate the mechanical response to force on integrins. Nature cell biology. 13 (6), 722–727. doi:10.1038/ncb2254.

Hama, T. & Park, F. (2016) Heterotrimeric G protein signaling in polycystic kidney disease. Physiological genomics. 48 (7), 429–445. doi:10.1152/physiolgenomics.00027.2016.

Happé, H., van der Wal, A.M., Leonhard, W.N., Kunnen, S.J., Breuning, M.H., de Heer, E. & Peters, D.J.M. (2011) Altered Hippo signalling in polycystic kidney disease. The Journal of pathology. 224 (1), 133–142. doi:10.1002/path.2856.

Hassane, S., Leonhard, W.N., van der Wal, A., Hawinkels, L.J., Lantinga-van Leeuwen, I.S., ten Dijke, P., Breuning, M.H., de Heer, E. & Peters, D.J. (2010) Elevated TGFbeta-Smad signalling in experimental Pkd1 models and human patients with polycystic kidney disease. The Journal of pathology. 222 (1), 21–31. doi:10.1002/path.2734.

Hayashi, K., Murai, T., Oikawa, H., Masuda, T., Kimura, K., Muehlich, S., Prywes, R. & Morita, T. (2015) A novel inhibitory mechanism of MRTF-A/B on the ICAM-1 gene expression in vascular endothelial cells. Scientific reports. 5, 10627. doi:10.1038/srep10627.

Hopp, K., Ward, C.J., Hommerding, C.J., Nasr, S.H., Tuan, H.-F., Gainullin, V.G., Rossetti, S., Torres, V.E. & Harris, P.C. (2012) Functional polycystin-1 dosage governs autosomal dominant polycystic kidney disease severity. The Journal of clinical investigation. 122 (11), 4257–4273. doi:10.1172/JCI64313.

Iwanciw, D., Rehm, M., Porst, M. & Goppelt-Struebe, M. (2003) Induction of connective tissue growth factor by angiotensin II: integration of signaling pathways. Arteriosclerosis, thrombosis, and vascular biology. 23 (10), 1782–1787. doi:10.1161/01.ATV.0000092913.60428.E6.

Kakiashvili, E., Speight, P., Waheed, F., Seth, R., Lodyga, M., Tanimura, S., Kohno, M., Rotstein, O.D., Kapus, A. & Szászi, K. (2009) GEF-H1 mediates tumor necrosis factor-alpha-induced Rho activation and myosin phosphorylation: role in the regulation of tubular paracellular permeability. The Journal of biological chemistry. 284 (17), 11454–11466. doi:10.1074/jbc.M805933200.

Kang, H.M., Ahn, S.H., Choi, P., Ko, Y.-A., Han, S.H., Chinga, F., Park, A.S.D., Tao, J., Sharma, K., Pullman, J., Bottinger, E.P., Goldberg, I.J. & Susztak, K. (2015) Defective fatty acid oxidation in renal tubular epithelial cells has a key role in kidney fibrosis development. Nature medicine. 21 (1), 37–46. doi:10.1038/nm.3762.

Kasahara, K. & Inagaki, M. (2021) Primary ciliary signaling: links with the cell cycle. Trends in cell biology. 31 (12), 954–964. doi:10.1016/j.tcb.2021.07.009.

Kim, T., Hwang, D., Lee, D., Kim, J.-H., Kim, S.-Y. & Lim, D.-S. (2017) MRTF potentiates TEAD-YAP transcriptional activity causing metastasis. The EMBO journal. 36 (4), 520–535. doi:10.15252/embj.201695137.

Kofler, M. & Kapus, A. (2021) Nucleocytoplasmic Shuttling of the Mechanosensitive Transcription Factors MRTF and YAP /TAZ. *Methods in molecular biology (Clifton*, N.J.). 2299, 197–216. doi:10.1007/978-1-0716-1382-5_15.

Kuwahara, K., Barrientos, T., Pipes, G.C.T., Li, S. & Olson, E.N. (2005) Muscle-specific signaling mechanism that links actin dynamics to serum response factor. Molecular and cellular biology. 25 (8), 3173–3181. doi:10.1128/MCB.25.8.3173-3181.2005.

Lakhia, R., Mishra, A., Biggers, L., Malladi, V., Cobo-Stark, P., Hajarnis, S. & Patel, V. (2023) Enhancer and super-enhancer landscape in polycystic kidney disease. Kidney international. 103 (1), 87–99. doi:10.1016/j.kint.2022.08.039.

Lanktree, M.B., Haghighi, A., di Bari, I., Song, X. & Pei, Y. (2021) Insights into Autosomal Dominant Polycystic Kidney Disease from Genetic Studies. Clinical journal of the American Society of Nephrology : CJASN. 16 (5), 790–799. doi:10.2215/CJN.02320220.

Leask, A. (2013) Integrin β1: A Mechanosignaling Sensor Essential for Connective Tissue Deposition by Fibroblasts. Advances in wound care. 2 (4), 160–166. doi:10.1089/wound.2012.0365.

Lee, H.S. (2012) Paracrine role for TGF-β-induced CTGF and VEGF in mesangial matrix expansion in progressive glomerular disease. Histology and histopathology. 27 (9), 1131–1141. doi:10.14670/HH-27.1131.

Lee, K., Boctor, S., Barisoni, L.M.C. & Gusella, G.L. (2015) Inactivation of integrin-β1 prevents the development of polycystic kidney disease after the loss of polycystin-1. Journal of the American Society of Nephrology : JASN. 26 (4), 888–895. doi:10.1681/ASN.2013111179.

Levy, M. & Feingold, J. (2000) Estimating prevalence in single-gene kidney diseases progressing to renal failure. Kidney international. 58 (3), 925–943. doi:10.1046/j.1523-1755.2000.00250.x.

Liu, C.-Y., Chan, S.W., Guo, F., Toloczko, A., Cui, L. & Hong, W. (2016) MRTF/SRF dependent transcriptional regulation of TAZ in breast cancer cells. Oncotarget. 7 (12), 13706–13716. doi:10.18632/oncotarget.7333.

Liu, X., Miao, J., Wang, C., Zhou, S., Chen, S., Ren, Q., Hong, X., Wang, Y., Hou, F.F., Zhou, L. & Liu, Y. (2020) Tubule-derived exosomes play a central role in fibroblast activation and kidney fibrosis. Kidney international. 97 (6), 1181–1195. doi:10.1016/j.kint.2019.11.026.

Lovisa, S., LeBleu, V.S., Tampe, B., Sugimoto, H., Vadnagara, K., Carstens, J.L., Wu, C.-C., Hagos, Y., Burckhardt, B.C., Pentcheva-Hoang, T., Nischal, H., Allison, J.P., Zeisberg, M. & Kalluri, R. (2015) Epithelial-to-mesenchymal transition induces cell cycle arrest and parenchymal damage in renal fibrosis. Nature medicine. 21 (9), 998–1009. doi:10.1038/nm.3902.

Luchsinger, L.L., Patenaude, C.A., Smith, B.D. & Layne, M.D. (2011) Myocardin-related transcription factor-A complexes activate type I collagen expression in lung fibroblasts. The Journal of biological chemistry. 286 (51), 44116–44125. doi:10.1074/jbc.M111.276931.

Luo, L., Roy, S., Li, L. & Ma, M. (2023) Polycystic kidney disease: novel insights into polycystin function. Trends in molecular medicine. 29 (4), 268–281. doi:10.1016/j.molmed.2023.01.005.

Mao, L., Liu, L., Zhang, T., Wu, X., Zhang, T. & Xu, Y. (2020) MKL1 mediates TGF-β-induced CTGF transcription to promote renal fibrosis. Journal of cellular physiology. 235 (5), 4790–4803. doi:10.1002/jcp.29356.

Márquez-Nogueras, K.M., Vuchkovska, V. & Kuo, I.Y. (2023) Calcium signaling in polycystic kidney disease-cell death and survival. Cell calcium. 112, 102733. doi:10.1016/j.ceca.2023.102733.

Masszi, A., Speight, P., Charbonney, E., Lodyga, M., Nakano, H., Szászi, K. & Kapus, A. (2010) Fate-determining mechanisms in epithelial-myofibroblast transition: major inhibitory role for Smad3. The Journal of cell biology. 188 (3), 383–399. doi:10.1083/jcb.200906155.

Menezes, L.F. & Germino, G.G. (2019) The pathobiology of polycystic kidney disease from a metabolic viewpoint. Nature reviews. Nephrology. 15 (12), 735–749. doi:10.1038/s41581-019-0183-y.

Miralles, F., Posern, G., Zaromytidou, A.-I. & Treisman, R. (2003) Actin dynamics control SRF activity by regulation of its coactivator MAL. Cell. 113 (3), 329–342. doi:10.1016/s0092-8674(03)00278-2.

Miranda, M.Z., Bialik, J.F., Speight, P., Dan, Q., Yeung, T., Szászi, K., Pedersen, S.F. & Kapus, A. (2017) TGF-β1 regulates the expression and transcriptional activity of TAZ protein via a Smad3-independent, myocardin-related transcription factor-mediated mechanism. The Journal of biological chemistry. 292 (36), 14902–14920. doi:10.1074/jbc.M117.780502.

Miranda, M.Z., Lichner, Z., Szászi, K. & Kapus, A. (2021) MRTF: Basic Biology and Role in Kidney Disease. International journal of molecular sciences. 22 (11). doi:10.3390/ijms22116040.

Morita, T., Mayanagi, T. & Sobue, K. (2007) Dual roles of myocardin-related transcription factors in epithelial mesenchymal transition via slug induction and actin remodeling. The Journal of cell biology. 179 (5), 1027–1042. doi:10.1083/jcb.200708174.

Muehlich, S., Rehm, M., Ebenau, A. & Goppelt-Struebe, M. (2017) Synergistic induction of CTGF by cytochalasin D and TGFβ-1 in primary human renal epithelial cells: Role of transcriptional regulators MKL1, YAP/TAZ and Smad2/3. Cellular signalling. 29, 31–40. doi:10.1016/j.cellsig.2016.10.002.

Ni, H., Li, A., Simonsen, N. & Wilkins, J.A. (1998) Integrin activation by dithiothreitol or Mn2+ induces a ligand-occupied conformation and exposure of a novel NH2-terminal regulatory site on the beta1 integrin chain. The Journal of biological chemistry. 273 (14), 7981–7987. doi:10.1074/jbc.273.14.7981.

Okada, H., Kikuta, T., Kobayashi, T., Inoue, T., Kanno, Y., Takigawa, M., Sugaya, T., Kopp, J.B. & Suzuki, H. (2005) Connective tissue growth factor expressed in tubular epithelium plays a pivotal role in renal fibrogenesis. Journal of the American Society of Nephrology : JASN. 16 (1), 133–143. doi:10.1681/ASN.2004040339.

Onuchic, L., Padovano, V., Schena, G., Rajendran, V., Dong, K., Shi, X., Pandya, R., Rai, V., Gresko, N.P., Ahmed, O., Lam, T.T., Wang, W., Shen, H., Somlo, S. & Caplan, M.J. (2023) The C-terminal tail of polycystin-1 suppresses cystic disease in a mitochondrial enzyme-dependent fashion. Nature communications. 14 (1), 1790. doi:10.1038/s41467-023-37449-1.

Orecchia, P., Conte, R., Balza, E., Castellani, P., Borsi, L., Zardi, L., Mingari, M.C. & Carnemolla, B. (2011) Identification of a novel cell binding site of periostin involved in tumour growth. European journal of cancer (Oxford, England : 1990). 47 (14), 2221–2229. doi:10.1016/j.ejca.2011.04.026.

Padovano, V., Podrini, C., Boletta, A. & Caplan, M.J. (2018) Metabolism and mitochondria in polycystic kidney disease research and therapy. Nature reviews. Nephrology. 14 (11), 678–687. doi:10.1038/s41581-018-0051-1.

Parnell, S.C., Magenheimer, B.S., Maser, R.L., Rankin, C.A., Smine, A., Okamoto, T. & Calvet, J.P. (1998) The polycystic kidney disease-1 protein, polycystin-1, binds and activates heterotrimeric G-proteins in vitro. Biochemical and biophysical research communications. 251 (2), 625–631. doi:10.1006/bbrc.1998.9514.

Podrini, C., Cassina, L. & Boletta, A. (2020) Metabolic reprogramming and the role of mitochondria in polycystic kidney disease. Cellular signalling. 67, 109495. doi:10.1016/j.cellsig.2019.109495.

Qiu, C., Huang, S., Park, J., Park, Y., Ko, Y.-A., Seasock, M.J., Bryer, J.S., Xu, X.-X., Song, W.-C., Palmer, M., Hill, J., Guarnieri, P., Hawkins, J., Boustany-Kari, C.M., Pullen, S.S., Brown, C.D. & Susztak, K. (2018) Renal compartment-specific genetic variation analyses identify new pathways in chronic kidney disease. Nature medicine. 24 (11), 1721–1731. doi:10.1038/s41591-018-0194-4.

Raman, A., Parnell, S.C., Zhang, Y., Reif, G.A., Dai, Y., Khanna, A., Daniel, E., White, C., Vivian, J.L. & Wallace, D.P. (2018) Periostin overexpression in collecting ducts accelerates renal cyst growth and fibrosis in polycystic kidney disease. American journal of physiology. Renal physiology. 315 (6), F1695– F1707. doi:10.1152/ajprenal.00246.2018.

Raman, A., Reif, G.A., Dai, Y., Khanna, A., Li, X., Astleford, L., Parnell, S.C., Calvet, J.P. & Wallace, D.P. (2017) Integrin-Linked Kinase Signaling Promotes Cyst Growth and Fibrosis in Polycystic Kidney Disease. Journal of the American Society of Nephrology : JASN. 28 (9), 2708–2719. doi:10.1681/ASN.2016111235.

Rayego-Mateos, S., Campillo, S., Rodrigues-Diez, R.R., Tejera-Muñoz, A., Marquez-Exposito, L., Goldschmeding, R., Rodríguez-Puyol, D., Calleros, L. & Ruiz-Ortega, M. (2021) Interplay between extracellular matrix components and cellular and molecular mechanisms in kidney fibrosis. Clinical science (London, England : 1979). 135 (16), 1999–2029. doi:10.1042/CS20201016.

Rozycki, M., Bialik, J.F., Speight, P., Dan, Q., Knudsen, T.E.T., Szeto, S.G., Yuen, D.A., Szászi, K., Pedersen, S.F. & Kapus, A. (2016) Myocardin-related Transcription Factor Regulates Nox4 Protein Expression: LINKING CYTOSKELETAL ORGANIZATION TO REDOX STATE. The Journal of biological chemistry. 291 (1), 227–243. doi:10.1074/jbc.M115.674606.

Sakai, N., Chun, J., Duffield, J.S., Wada, T., Luster, A.D. & Tager, A.M. (2013) LPA1-induced cytoskeleton reorganization drives fibrosis through CTGF-dependent fibroblast proliferation. FASEB journal : official publication of the Federation of American Societies for Experimental Biology. 27 (5), 1830–1846. doi:10.1096/fj.12-219378.

Schuster, R., Younesi, F., Ezzo, M. & Hinz, B. (2023) The Role of Myofibroblasts in Physiological and Pathological Tissue Repair. Cold Spring Harbor perspectives in biology. 15 (1). doi:10.1101/cshperspect.a041231.

Shen, B., Estevez, B., Xu, Z., Kreutz, B., Karginov, A., Bai, Y., Qian, F., Norifumi, U., Mosher, D. & Du, X. (2015) The interaction of Gα13 with integrin β1 mediates cell migration by dynamic regulation of RhoA. Molecular biology of the cell. 26 (20), 3658–3670. doi:10.1091/mbc.E15-05-0274.

Small, E.M., Thatcher, J.E., Sutherland, L.B., Kinoshita, H., Gerard, R.D., Richardson, J.A., Dimaio, J.M., Sadek, H., Kuwahara, K. & Olson, E.N. (2010) Myocardin-related transcription factor-a controls myofibroblast activation and fibrosis in response to myocardial infarction. Circulation research. 107 (2), 294–304. doi:10.1161/CIRCRESAHA.110.223172.

Song, X., Di Giovanni, V., He, N., Wang, K., Ingram, A., Rosenblum, N.D. & Pei, Y. (2009a) Systems biology of autosomal dominant polycystic kidney disease (ADPKD): computational identification of gene expression pathways and integrated regulatory networks. Human molecular genetics. 18 (13), 2328– 2343. doi:10.1093/hmg/ddp165.

Song, X., Di Giovanni, V., He, N., Wang, K., Ingram, A., Rosenblum, N.D. & Pei, Y. (2009b) Systems biology of autosomal dominant polycystic kidney disease (ADPKD): computational identification of gene expression pathways and integrated regulatory networks. Human molecular genetics. 18 (13), 2328– 2343. doi:10.1093/hmg/ddp165.

Speight, P., Kofler, M., Szászi, K. & Kapus, A. (2016) Context-dependent switch in chemo/mechanotransduction via multilevel crosstalk among cytoskeleton-regulated MRTF and TAZ and TGFβ-regulated Smad3. Nature communications. 7, 11642. doi:10.1038/ncomms11642.

Speight, P., Nakano, H., Kelley, T.J., Hinz, B. & Kapus, A. (2013) Differential topical susceptibility to TGFβ in intact and injured regions of the epithelium: key role in myofibroblast transition. Molecular biology of the cell. 24 (21), 3326–3336. doi:10.1091/mbc.E13-04-0220.

Streets, A.J., Prosseda, P.P. & Ong, A.C. (2020) Polycystin-1 regulates ARHGAP35-dependent centrosomal RhoA activation and ROCK signaling. JCI insight. 5 (16). doi:10.1172/jci.insight.135385.

Suzuki, N., Hajicek, N. & Kozasa, T. (2009) Regulation and physiological functions of G12/13-mediated signaling pathways. Neuro-Signals. 17 (1), 55–70. doi:10.1159/000186690.

Suzuki, N., Nakamura, S., Mano, H. & Kozasa, T. (2003) Galpha 12 activates Rho GTPase through tyrosine-phosphorylated leukemia-associated RhoGEF. Proceedings of the National Academy of Sciences of the United States of America. 100 (2), 733–738. doi:10.1073/pnas.0234057100.

Szeto, S.G., Narimatsu, M., Lu, M., He, X., Sidiqi, A.M., Tolosa, M.F., Chan, L., De Freitas, K., Bialik, J.F., Majumder, S., Boo, S., Hinz, B., Dan, Q., Advani, A., John, R., Wrana, J.L., Kapus, A. & Yuen, D.A. (2016) YAP/TAZ Are Mechanoregulators of TGF-β-Smad Signaling and Renal Fibrogenesis. Journal of the American Society of Nephrology : JASN. 27 (10), 3117–3128. doi:10.1681/ASN.2015050499.

Tan, R.J., Zhou, D. & Liu, Y. (2016) Signaling Crosstalk between Tubular Epithelial Cells and Interstitial Fibroblasts after Kidney Injury. *Kidney diseases (Basel*, Switzerland). 2 (3), 136–144. doi:10.1159/000446336.

Tanner, G.A., McQuillan, P.F., Maxwell, M.R., Keck, J.K. & McAteer, J.A. (1995) An in vitro test of the cell stretch-proliferation hypothesis of renal cyst enlargement. Journal of the American Society of Nephrology : JASN. 6 (4), 1230–1241. doi:10.1681/ASN.V641230.

Traykova-Brauch, M., Schönig, K., Greiner, O., Miloud, T., Jauch, A., Bode, M., Felsher, D.W., Glick, A.B., Kwiatkowski, D.J., Bujard, H., Horst, J., von Knebel Doeberitz, M., Niggli, F.K., Kriz, W., Gröne, H.-J. & Koesters, R. (2008) An efficient and versatile system for acute and chronic modulation of renal tubular function in transgenic mice. Nature medicine. 14 (9), 979–984. doi:10.1038/nm.1865.

Vartiainen, M.K., Guettler, S., Larijani, B. & Treisman, R. (2007) Nuclear actin regulates dynamic subcellular localization and activity of the SRF cofactor MAL. *Science (New York*, N.Y.). 316 (5832), 1749– 1752. doi:10.1126/science.1141084.

Waheed, F., Dan, Q., Amoozadeh, Y., Zhang, Y., Tanimura, S., Speight, P., Kapus, A. & Szászi, K. (2013) Central role of the exchange factor GEF-H1 in TNF-α-induced sequential activation of Rac, ADAM17/TACE, and RhoA in tubular epithelial cells. Molecular biology of the cell. 24 (7), 1068–1082. doi:10.1091/mbc.E12-09-0661.

Wallace, D.P., White, C., Savinkova, L., Nivens, E., Reif, G.A., Pinto, C.S., Raman, A., Parnell, S.C., Conway, S.J. & Fields, T.A. (2014) Periostin promotes renal cyst growth and interstitial fibrosis in polycystic kidney disease. Kidney international. 85 (4), 845–854. doi:10.1038/ki.2013.488.

Wang, D., Prakash, J., Nguyen, P., Davis-Dusenbery, B.N., Hill, N.S., Layne, M.D., Hata, A. & Lagna, G. (2012a) Bone morphogenetic protein signaling in vascular disease: anti-inflammatory action through myocardin-related transcription factor A. The Journal of biological chemistry. 287 (33), 28067–28077. doi:10.1074/jbc.M112.379487.

Wang, D., Prakash, J., Nguyen, P., Davis-Dusenbery, B.N., Hill, N.S., Layne, M.D., Hata, A. & Lagna, G. (2012b) Bone morphogenetic protein signaling in vascular disease: anti-inflammatory action through myocardin-related transcription factor A. The Journal of biological chemistry. 287 (33), 28067–28077. doi:10.1074/jbc.M112.379487.

Wang, L. & Dynlacht, B.D. (2018) The regulation of cilium assembly and disassembly in development and disease. *Development (Cambridge*, England). 145 (18). doi:10.1242/dev.151407.

Wipff, P.-J., Majd, H., Acharya, C., Buscemi, L., Meister, J.-J. & Hinz, B. (2009) The covalent attachment of adhesion molecules to silicone membranes for cell stretching applications. Biomaterials. 30 (9), 1781– 1789. doi:10.1016/j.biomaterials.2008.12.022.

Wu, G., D’Agati, V., Cai, Y., Markowitz, G., Park, J.H., Reynolds, D.M., Maeda, Y., Le, T.C., Hou, H., Kucherlapati, R., Edelmann, W. & Somlo, S. (1998) Somatic inactivation of Pkd2 results in polycystic kidney disease. Cell. 93 (2), 177–188. doi:10.1016/s0092-8674(00)81570-6.

Wu, Y., Xu, J.X., El-Jouni, W., Lu, T., Li, S., Wang, Q., Tran, M., Yu, W., Wu, M., Barrera, I.E., Bonventre, J. V, Zhou, J., Denker, B.M. & Kong, T. (2016) Gα12 is required for renal cystogenesis induced by Pkd1 inactivation. Journal of cell science. 129 (19), 3675–3684. doi:10.1242/jcs.190496.

Xu, H., Wu, X., Qin, H., Tian, W., Chen, J., Sun, L., Fang, M. & Xu, Y. (2015) Myocardin-Related Transcription Factor A Epigenetically Regulates Renal Fibrosis in Diabetic Nephropathy. Journal of the American Society of Nephrology : JASN. 26 (7), 1648–1660. doi:10.1681/ASN.2014070678.

Yamamura, Y., Sakai, N., Iwata, Y., Lagares, D., Hara, A., et al. (2023) Myocardin-related transcription factor contributes to renal fibrosis through the regulation of extracellular microenvironment surrounding fibroblasts. FASEB journal : official publication of the Federation of American Societies for Experimental Biology. 37 (7), e23005. doi:10.1096/fj.202201870R.

Yang, Q., Mcgowen, K.M., Yin, J., Alilin, A.N., Karzai, F.H., Dahut, W.L. & Corey, E. (2018) *Downloaded from clincancerres.aacrjournals.org on June 25, 2018. © 2018 American Association for Cancer Research*. doi:10.1158/1078-0432.CCR-18-0409.

Yao, L., Zhou, Y., Li, J., Wickens, L., Conforti, F., et al. (2021) Bidirectional epithelial-mesenchymal crosstalk provides self-sustaining profibrotic signals in pulmonary fibrosis. The Journal of biological chemistry. 297 (3), 101096. doi:10.1016/j.jbc.2021.101096.

Yin, Q. & Liu, H. (2019) Connective Tissue Growth Factor and Renal Fibrosis. Advances in experimental medicine and biology. 1165, 365–380. doi:10.1007/978-981-13-8871-2_17.

Yook, Y.J., Woo, Y.M., Yang, M.H., Ko, J.Y., Kim, B.H., Lee, E.J., Chang, E.S., Lee, M.J., Lee, S. & Park, J.H. (2012) Differential Expression of PKD2-Associated Genes in Autosomal Dominant Polycystic Kidney Disease. Genomics & informatics. 10 (1), 16–22. doi:10.5808/GI.2012.10.1.16.

Yu, O.M., Miyamoto, S. & Brown, J.H. (2016) Myocardin-Related Transcription Factor A and Yes-Associated Protein Exert Dual Control in G Protein-Coupled Receptor- and RhoA-Mediated Transcriptional Regulation and Cell Proliferation. Molecular and cellular biology. 36 (1), 39–49. doi:10.1128/MCB.00772-15.

Yuasa, T., Takakura, A., Denker, B.M., Venugopal, B. & Zhou, J. (2004) Polycystin-1L2 is a novel G-protein-binding protein. Genomics. 84 (1), 126–138. doi:10.1016/j.ygeno.2004.02.008.

Zhang, Y., Dai, Y., Raman, A., Daniel, E., Metcalf, J., Reif, G., Pierucci-Alves, F. & Wallace, D.P. (2020) Overexpression of TGF-β1 induces renal fibrosis and accelerates the decline in kidney function in polycystic kidney disease. American journal of physiology. Renal physiology. 319 (6), F1135–F1148. doi:10.1152/ajprenal.00366.2020.

Zhang, Y., Reif, G. & Wallace, D.P. (2020) Extracellular matrix, integrins, and focal adhesion signaling in polycystic kidney disease. Cellular signalling. 72, 109646. doi:10.1016/j.cellsig.2020.109646.

Zhou, D., Fu, H., Zhang, L., Zhang, K., Min, Y., Xiao, L., Lin, L., Bastacky, S.I. & Liu, Y. (2017) Tubule-Derived Wnts Are Required for Fibroblast Activation and Kidney Fibrosis. Journal of the American Society of Nephrology : JASN. 28 (8), 2322–2336. doi:10.1681/ASN.2016080902.

## Supplementary References

1. Speight, P., Kofler, M., Szászi, K. & Kapus, A. Context-dependent switch in chemo/mechanotransduction via multilevel crosstalk among cytoskeleton-regulated MRTF and TAZ and TGFβ-regulated Smad3. Nat Commun 7, 11642 (2016).

2. Miranda, M. Z. et al. TGF-β1 regulates the expression and transcriptional activity of TAZ protein via a Smad3-independent, myocardin-related transcription factor-mediated mechanism. J Biol Chem 292, 14902–14920 (2017).

3. Kofler, M. & Kapus, A. Nucleocytoplasmic Shuttling of the Mechanosensitive Transcription Factors MRTF and YAP /TAZ. Methods Mol Biol 2299, 197–216 (2021).

4. Goffin, J. M. et al. Focal adhesion size controls tension-dependent recruitment of alpha-smooth muscle actin to stress fibers. J Cell Biol 172, 259–68 (2006).

5. Wipff, P.-J. et al. The covalent attachment of adhesion molecules to silicone membranes for cell stretching applications. Biomaterials 30, 1781–9 (2009).

6. Bialik, J. F. et al. Profibrotic epithelial phenotype: a central role for MRTF and TAZ. Sci Rep 9, 4323 (2019).

7. Szeto SG, Narimatsu M, Lu M, He X, Sidiqi AM, Tolosa MF, Chan L, De Freitas K, Bialik JF, Majumder S, Boo, Hinz B, Dan Q, Advani A, John R, Wrana JL, Kapus A, Y. DA. YAP/TAZ Are Mechanoregulators of TGF-β-Smad Signaling and Renal Fibrogenesis. Journal of American Society of NephrologyAmerican Society of 27, 3117–3128 (2016).

8. Patro, R., Duggal, G., Love, M. I., Irizarry, R. A. & Kingsford, C. Salmon provides fast and bias-aware quantification of transcript expression. Nat Methods 14, (2017).

9. Soneson, C., Love, M. I. & Robinson, M. D. Differential analyses for RNA-seq: Transcript-level estimates improve gene-level inferences. F1000Res 4, (2016).

10. Langfelder, P. & Horvath, S. WGCNA: An R package for weighted correlation network analysis. BMC Bioinformatics 9, (2008).

11. Love, M. I., Huber, W. & Anders, S. Moderated estimation of fold change and dispersion for RNA-seq data with DESeq2. Genome Biol 15, (2014).

12. Kim, D., Paggi, J. M., Park, C., Bennett, C. & Salzberg, S. L. Graph-based genome alignment and genotyping with HISAT2 and HISAT-genotype. Nat Biotechnol 37, (2019).

13. Ihaka, R. & Gentleman, R. R: A Language for Data Analysis and Graphics. Journal of Computational and Graphical Statistics 5, (1996).

14. Liao, Y., Smyth, G. K. & Shi, W. The R package Rsubread is easier, faster, cheaper and better for alignment and quantification of RNA sequencing reads. Nucleic Acids Res 47, (2019).

15. Anders, S., Reyes, A. & Huber, W. Detecting differential usage of exons from RNA-seq data. Genome Res 22, (2012).

16. Mootha, V. K. et al. PGC-1α-responsive genes involved in oxidative phosphorylation are coordinately downregulated in human diabetes. Nat Genet 34, (2003).

17. Subramanian, A. et al. Gene set enrichment analysis: A knowledge-based approach for interpreting genome-wide expression profiles. Proc Natl Acad Sci U S A 102, (2005).

18. Liberzon, A. et al. The Molecular Signatures Database Hallmark Gene Set Collection. Cell Syst 1, (2015).

19. Ge, S. X., Jung, D., Jung, D. & Yao, R. ShinyGO: A graphical gene-set enrichment tool for animals and plants. Bioinformatics 36, (2020).

